# HaploCharmer: a Snakemake workflow for read-scale haplotype calling adapted to polyploids

**DOI:** 10.1101/2025.03.14.642807

**Authors:** Simon Rio, Sophie Abdallah, Théo Durand, Angélique D’Hont, Olivier Garsmeur

## Abstract

The advent of next-generation sequencing (NGS) has revolutionized the study of single nucleotide polymorphisms (SNPs), making it increasingly cost-effective. Haplotypes, which combine alleles from adjacent variants, offer several advantages over bi-allelic SNPs, including enhanced information content, reduced dimensionality, and improved statistical power in genomic studies. These benefits are particularly significant for polyploid species, where distinguishing all homologous copies using SNP markers alone can be challenging. This article introduces HaploCharmer, a flexible workflow designed for read-scale haplotype calling from NGS data. HaploCharmer identifies haplotypes within preconfigured genomic regions smaller than a sequencing read, ensuring direct comparability across individuals. It integrates a series of processing steps including mapping, haplotype identification, filtration, and reporting of haplotype sequences, as presence-absence, in the panel of accessions analyzed. The performance of HaploCharmer was validated by building a genetic map using whole-genome sequencing data from a highly polyploid sugarcane cultivar (R570) and its self-progeny, and performing a diversity analysis in the polyploid *Saccharum* genus using targeted sequencing data. The workflow successfully identified a large number of high-quality haplotypes, with less than 1% of false positives. The dense genetic map obtained using single-dose haplotypes accurately depicted the known genome architecture of the R570 cultivar, including large chromosome rearrangements. The diversity analysis accurately reflected the known genetic structure within this genus. It also allowed inferring ancestral origins to mapped haplotypes and the corresponding chromosome segments in the R570 genetic map. HaploCharmer provides a robust method for diversity, genetic mapping, and quantitative genetics studies in both diploid and polyploid species.

## Introduction

Because of their convenience in subsequent analyses, most variant call methods from next-generation sequencing (NGS) data aim to characterize bi-allelic single nucleotide polymorphisms (SNPs). Alternatively, alleles from adjacent variants, including SNPs and other polymorphisms such as indels, can be combined into haplotypes. The advantages of haplotypes over bi-allelic SNPs include an increased information content and reduced dimensionality, allowing better capture of linkage disequilibrium, higher statistical power in genome-wide association studies, higher genomic prediction accuracy, better genetic mapping resolution and improved population structure inference (Bhat et al., 2021; Gattepaille and Jakobsson, 2012). The multi-allelic nature of haplotypic markers is expected to be particularly useful in polyploid species, where all homologous copies cannot be distinguished by a single bi-allelic variant, unlike diploid species (Voorrips and Tumino, 2022).

Using NGS data, alleles from adjacent variants can be combined into haplotypes when they are observed on the same sequencing read. This strategy is implemented in HAPLOSWEEP (Clevenger et al., 2018), but is limited to phasing pairs of variants for polyploid species. In addition, haplotypes can be extended by assembling reads on the basis of their shared allele content, which supports an origin from the same homologous copy. This task is more complex in polyploids for which the solution space increases dramatically with the number of variants and the ploidy level (He et al., 2018). Several haplotype assembling methods have been proposed in the literature, some of them specifically dedicated to polyploids, such as HapCompass (Aguiar and Istrail, 2012, 2013), Poly-Harsh (He et al., 2018), or Ranbow (Moeinzadeh et al., 2020). However, these approaches can lead to considerable heterogeneity in haplotype length between homologous copies of the same individual or between different individuals, depending on the degree of information available to guide assembly. As a consequence, the resulting haplotypes may not be easily transformed into multi-allelic markers to be used in genomic and genetic studies.

In this article, we present HaploCharmer, a workflow for read-scale haplotype calling implemented in Snakemake (Köster and Rahmann, 2012; Mölder et al., 2021). Haplotypes are called within preconfigured genomic regions of a size smaller than a sequencing read, and within which all possible variants are considered: bi- and multi-allelic SNPs as well as small indels. Like in HAPLOSWEEP, alleles are considered as phased only if they are jointly observed on the same sequencing read. The use of preconfigured genomic regions, within which haplotypes are characterized, ensures that the resulting haplotypes are directly comparable across individuals. Sugarcane is a major crop for sugar and energy production, and has one of the most complex genomes of all cultivated plants.

Modern sugarcane cultivars are the result of relatively recent inter-specific hybridization (a century ago) between two polyploid species: the sugar-producing species *Saccharum officinarum* (2n=8x=80, x=10) and the wild species *Saccharum spontaneum* (2n=5x=40 to 16x=128, x=8). In *S. spontaneum*, two sets of three chromosomes have each been rearranged into two chromosomes, in comparison with the ancestral chromosome structure observed in *S. officinarum* (Garsmeur et al., 2018; Piperidis and D’Hont, 2020). Modern cultivars are thus complex polyploids, with on average twelve copies for each basic chromosome, of which 15 to 25% of the chromosomes are derived from *S. spontaneum*, including recombinant with *S. officinarum* (D’Hont et al., 1996; Piperidis et al., 2010; Piperidis and D’Hont, 2020).

HaploCharmer’s effectiveness in identifying haplotypes in polyploid and heterozygous genomes was assessed by building a genetic map from a self-progeny of a sugarcane cultivar (R570) genotyped for 87K genomic regions, and by analysing a diversity panel of 307 *Saccharum* accessions.

## Materials and methods

### HaploCharmer workflow

HaploCharmer takes as input single or paired-end short-read sequencing data (FASTQ) for a set of samples, which have passed quality controls such as adapter removal, read filtering and trimming of low-quality bases.

First, reads are mapped to a reference sequence genome (FASTA) using BWA-MEM (Li, 2013). Standard processing steps are applied to BAM files, such as removal of read duplicates using Pi-card MarkDuplicates (https://broadinstitute.github.io/picard/), local realignment around indels using GATK3 (DePristo et al., 2011), and update of the string encoding mismatched and deleted reference bases (MD) flag using SAMtools (Danecek et al., 2021). Haplotypes are then identified and counted within pre-configured genomic coordinates (BED) using a python script (BAM_to_gVCF.py) based on the pysam module (https://github.com/pysam-developers/pysam/). Only sequencing reads whose alignment completely covers the predefined genomic regions - also called phase sets (PSs) - are considered (Figure 1A). While PSs can vary in size, they must not overlap. For each of the samples analyzed, all positions that fall into a PS are reported in a genomic VCF file (gVCF), regardless of whether a variant (SNP or indel) was detected at that site or not. In the gVCF file, haplotypes are reported as groups of alleles phased within each PS, which is specified following the convention of using the pipe “|” in the genotype (GT) field. Individual sample files are merged into a global VCF file using BCFtools (Danecek et al., 2021), and filtered for haplotypes likely resulting from sequencing errors using a python script (VCF_filter.py). For this filtering step, the general approach is to discard haplotypes that are never observed with a high degree of certainty in at least one sample, i.e. with sufficient read depth and frequency to support the existence of the haplotype (see next subsection “HaploCharmer parameters”). Then, a non-standard file characterizing haplotype presence-absence (HPA) is generated using a python script (VCF_to_HapPresAbs.py) based on the pyfaidx module (Shirley et al., 2015). In this HPA file, each line corresponds to a haplotype and includes its coordinates on the reference sequence, its complete sequence, and sample-specific columns indicating its presence/absence genotyping along with read depth information (Figure 1B). In addition, the nature and position of variants (SNPs and indels) distinguishing the haplotypes of a same PS are reported in a text file (INFO). This file is made up of a concatenation of tables in VCF format, one for each PS, where haplotypes are represented in columns. Each sub-table corresponding to a specific PS can easily be accessed using a “grep” command with the name of the PS as the argument. Finally, a report (HTML) is generated presenting various statistics, such as read mapping, read duplicates, phase set read depths, and number of haplotypes per phase set. The workflow’s key steps are summarized in Figure 1C.

**Figure 1.**
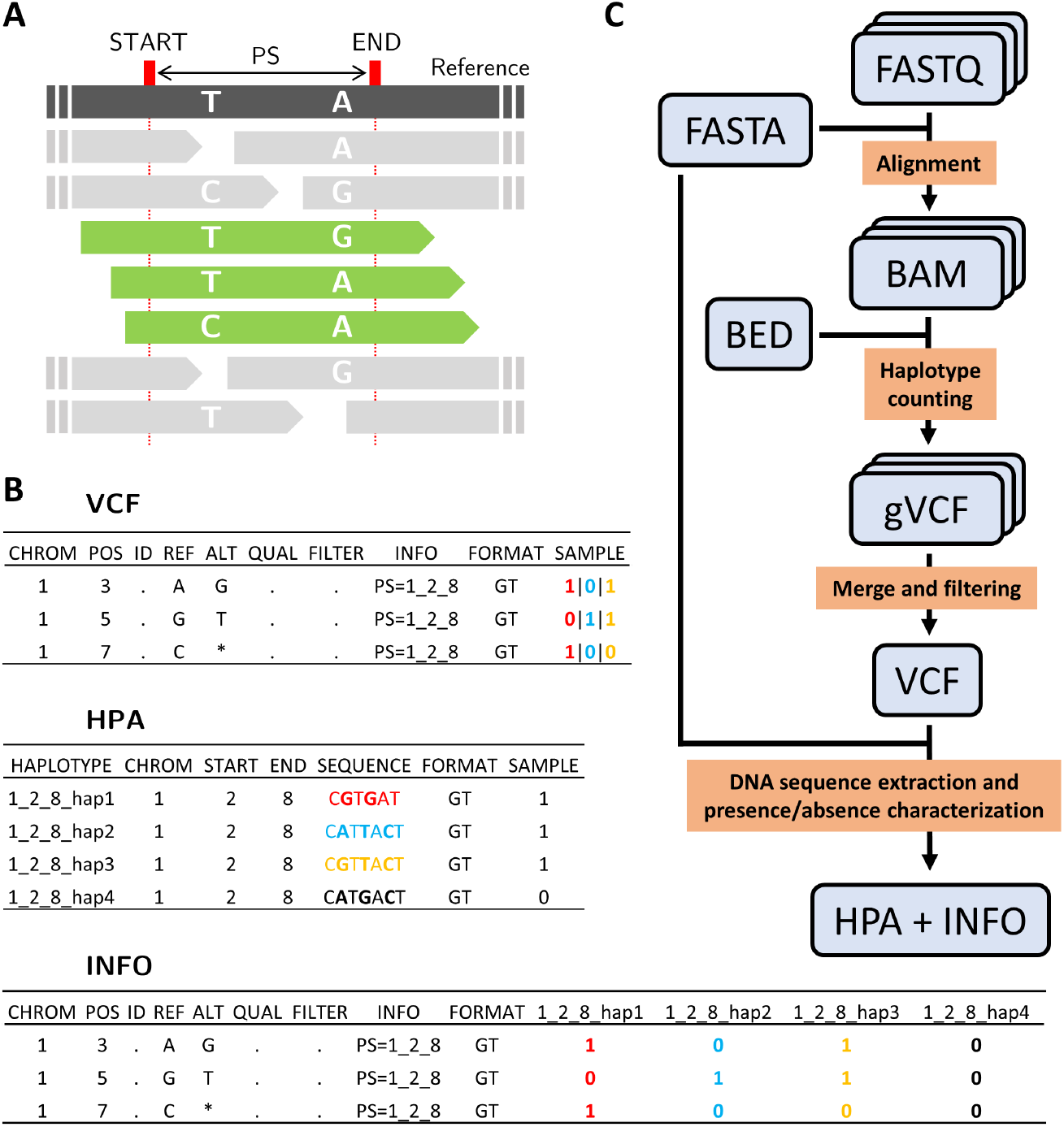
(A) Schematic representation of three aligned reads considered for haplotyping (green) that overlap the entire phase set (PS) defined by the START and END positions on the reference, unlike the other reads (grey). (B) Illustration of the haplotype representation in the VCF, HPA and INFO files for one sample genotyped for a genomic region “1_2_8”, i.e. a phase set spanning chromosome 1 from position 2 to 8. For the HPA and INFO files, the haplotype “1_2_8_hap4” corresponds to the reference sequence haplotype. Only the genotype (GT) format field is represented here although it is normally associated with read depth information in real files. (C) Summary of the HaploCharmer (i) inputs including the reference sequence (FASTA), sequencing reads (FASTQ), PS coordinates on the reference sequence (BED), (ii) intermediate files including alignment files (BAM) and genomic VCF files (gVCF), (iii) and outputs including VCF, HPA and INFO files. Key workflow steps to obtain output from input files are described in orange rectangles.

### HaploCharmer parameters

The haplotype calling relies on a limited set of parameters that are specified in the Snakemake YAML configuration file. Regarding the mapping, the removing of read duplicates is optional, although recommended, and a minimum mapping quality (i.e. MAPQ provided by BWA-MEM) must be specified. For the VCF filtering step, a genotyping data point must be supported by a read depth (DP) comprised between “min_gt_depth” (minimum read depth) and “max_gt_depth” (maximum read depth) otherwise the genotyping datapoint is set to a missing value. Then, each haplotype must be supported by a read depth (AD) greater than or equal to “min_hap_depth” and a frequency (AD/DP) greater than or equal to “min_hap_freq” in at least one sample. Otherwise, this haplotype is considered as likely resulting from sequencing errors and discarded from the set of haplotypes in the population.

Although a haplotype may be supported at a population level, it may be the result of sequencing errors for a particular sample. At the sample level, the haplotypes reported in the HPA file are first re-inspected to verify that they are supported by a read depth (AD) and frequency (AD/DP) superior or equal to “min_pres_depth” and “min_pres_freq”, respectively. Then, too rare haplotypes with a frequency below “max_abs_freq”, are considered absent in the sample. Finally, haplotypes that do not meet any of these criteria (e.g. intermediate frequency: “max_abs_freq” *≤* AD/DP *<* “min_pres_freq”) are set as missing values.

The workflow parameters are explained in further details and illustrated with a toy example in the README.md file of the Gitlab repository (see “Data, script, code, and supplementary information availability” section).

### Evaluation of HaploCharmer on polyploid sugarcane

The performance of HaploCharmer was assessed by calling haplotypes from Illumina wholegenome sequencing data from sugarcane cultivar R570 (*≈*12x) and 96 accessions of its self-progeny (Grivet et al., 1994; Grivet et al., 1996). The identified haplotypes were then characterized to select segregating single-dose haplotypes, which were used to construct a genetic map. In addition, a diversity panel of 307 *Saccharum* accessions and related genera, for which targeted enrichment sequencing data were publicly available (Yang et al., 2018), was characterized for a large part of the same genomic regions.

#### Haplotyping of R570 and its self-progeny

The recent polyploid genome assembly of R570 (Healey et al., 2024) was used as a sugarcane reference sequence. From this polyploid assembly, with several homologous copies assembled for each basic chromosome (i.e. homology group), a monoploid reference sequence was derived by choosing the longest scaffolds from each homology group, but excluding those representing rearranged chromosomes. The scaffolds considered were the following: Chr1A, Chr2A, Chr3A, Chr4A, Chr5A, Chr6A, Chr7A, Chr8A, Chr9A, and Chr10A.

Three sets of probes were used to define genomic regions: 60,000 120 bp probes defined by Yang et al. (2018), 1,964 120 bp probes (non-redundant with the previous set) from the set of 19,436 probes defined in Dijoux et al. (2024), and 39,867 additional 80 bp probes defined in sugarcane gene models. All probes were mapped onto the monoploid reference genome using BWA-MEM (Li, 2013). Different filters were applied, excluding probes that i) were not uniquely mapped, ii) mapped on an interval below 60 bp or above 140 bp, and iii) overlapped the mapping coordinates of another adjacent probe. A total of 86,681 80 bp genomic regions distributed along the R570 monoploid genome assembly were retained and were considered as PSs for the haplotype calling. The coordinates of these PSs were reported in a BED file format.

To reduce the computational time required for aligning high-depth WGS data from cultivar R570 and the 96 progenies, the reads corresponding to all regions were recovered through insilico capture using probe sequences as targets with Mirabait (Chevreux et al., 2004).

The HaploCharmer workflow was launched by specifying the following parameters in the Snakemake YAML configuration file:

- Remove_duplicates: True
- Mapping_quality: 20
- VCF_filter: “–min_gt_depth 30 –max_gt_depth 1000 –min_hap_depth 4 –min_hap_freq 0.04”
- VCF_to_HapPresAbs: “–max_abs_freq 0.01 –min_pres_depth 3 –min_pres_freq 0.04”

#### Filtering of progenies

From the raw haplotypes identified and reported in the HPA output file, progenies’ genotypes (present, absent, or missing) were compared to that obtained for R570 to potentially identify illegitimate progeny (Figure S1). No progeny was found to have a substantial proportion of haplotypes absent from R570, suggesting no illegitimate cross. However, for a self-progeny, a substantial proportion of haplotypes is expected to be present in the parent but absent in the progeny, notably resulting from the segregation of single-dose haplotypes (see subsection “Characterization of single-dose markers”). Few progenies were discarded from the study as they presented a genotyping profile too similar to R570, suggesting that there were clones of the parent: AF411, AF413, AF439, AF454, AF491, AF499, and AF568. Two additional progenies were discarded due to a significant percentage of missing values: AF450 and AF488. After filtering, a total of 87 progenies were conserved for downstream analyses.

#### Filtering of haplotypes

A total of 247,267 haplotypes distributed in 73,100 phase sets were obtained from the raw presence-absence HPA output file. Given the ploidy level of R570 (*≈*12x with limited aneuploidy), and the self-progeny nature of the population, a maximum of around 12 haplotypes can theoretically be expected from each PS. As a consequence, PSs with more than 14 haplotypes were discarded as they likely resulted from reads originating from duplicated regions. In addition, PSs revealing only one haplotype were discarded from the dataset, as well as haplotypes with more than 25% missing values. A total of 215,704 haplotypes distributed in 67,917 PSs were conserved for the 87 progenies. The number of haplotypes per PS varied from 2 to 14, with an average of 3.17.

#### SNPs and indels contributing to haplotypes

The ability of the haplotypes genotyped by HaploCharmer to reveal single-dose markers was compared to that of unphased variants (biallelic SNPs, multiallelic SNPs, indels) contributing those haplotypes, as reported in the VCF output file. The total 215,704 haplotypes resulted from the combination of 292,518 variants. The presence/absence genotyping of each allele of these variants was re-inspected using the same parameters used to generate the HPA file (see “HaploCharmer parameters” sub-section). To this end, allele read depths that are initially reported specifically for each haplotype were aggregated per allele. After filtering out alleles with more than 25% missing values 504,558 alleles were conserved: 352,240 alleles from bi-allelic SNPs (69.81%), 130,745 alleles from bi-allelic indels (25.91%), 6,700 alleles from multi-allelic SNPs (1.33%), and 14,873 other more complex variants (2.95%).

#### Characterization of single-dose markers

Single-dose markers (a single copy is present at the locus in the parent, unlike multi-dose markers) have a 50% chance of being transmitted to the gametes, resulting in an expected presence:absence ratio of 3:1 in a self-progeny. The probability of transmission of multi-dose markers depends on the number of doses, the ploidy and the pairing of homologous chromosomes during meiosis. For instance, assuming a ploidy of 12, as in sugarcane cultivars, and random bivalent chromosome pairing at meiosis (i.e. polysomic inheritance), double-dose markers have 77% chance of being transmitted to the gametes. In presence of strict preferential bivalent pairing of chromosomes (i.e. disomic inheritance), doubledose markers have a 75% or 100% chance of being transmitted to gametes if the two doses are located on unpaired or paired chromosomes, respectively. As a consequence, the expected presence:absence ratio of multi-dose markers is always higher than 3:1 in the self-progeny of a polyploid species like sugarcane.

To classify *M* markers (haplotypes, SNPs or indels) based on their segregation in the self-progeny population, a mixture model was proposed in which the latent type *Z*_*m*_ of marker *m* is drawn from a Multinomial distribution:

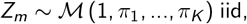

where *π*_*k*_ is the proportion of markers from type *k* (false positives, single-dose or multi-dose) and iid stands for independent and identically distributed. The number of progenies *X*_*m*_ genotyped as present for the marker is drawn from a Binomial distribution conditional on the marker type:

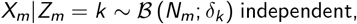

where *N*_*m*_ is the number of progenies genotyped for the marker (excluding missing values), and *δ*_*k*_ is the expected proportion of progenies genotyped as present. The inference of model parameters is done using an expectation-maximization (EM) algorithm (Dempster et al., 1977). In the expectation step, the conditional probability ℙ (*Z*_*m*_ = *k*|*X*_*m*_) = *τ*_*mk*_ is calculated:

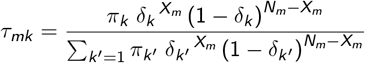

In the maximization step, the values of parameters *π*_*k*_ and *δ*_*k*_ are updated as following:

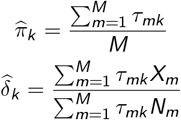

Parameters were initialized as 1/3 for all *π*_*k*_, and *δ*_*k*_ as 0.1, 0.75, and 0.9 for putative false positives, single-dose and multi-dose markers categories, respectively. The EM algorithm iterates until convergence, i.e. a difference of less than 0.001 between two iterations, of the marginal log-likelihood:

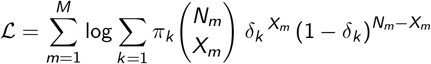

Haplotypes and other markers were considered single-dose in R570 when the probability of being a single dose marker (obtained from *τ*_*mk*_) was at least 0.99.

#### Genetic map

Single-dose haplotypes were grouped into non-redundant bins used as effective markers for genetic mapping using the R-package “onemap” (Margarido et al., 2007). After estimating the recombination fraction between all bin pairs, haplotype bins were clustered into cosegregation groups considering a LOD score of 7. Each cosegregation group was assigned to a basic chromosome (i.e. homology group) based on the mapping position of the majority of haplotypes from the group. Note that known rearranged chromosomes in R570 (i.e. chromosomes 8/5, 5/9, 6/9, 7/10 and 8/10) were also taken into account as they displayed large numbers of haplotypes located on two chromosomes of the monoploid reference (Healey et al., 2024). Haplotypes whose position on the reference were discordant with the chromosome class or the recombination fraction with neighboring haplotypes on the chromosome were filtered out.

Single-dose markers corresponding to SNPs and indels contributing to haplotypes were also grouped into non-redundant bins to compare the number of effective markers obtained.

#### Haplotyping of the diversity panel

A diversity panel of 307 accessions of *Saccharum* species (*S. spontaneum, S. robustum, S. officinarum, S. barberi, S. sinense* and hybrids) and related genera, for which targeted enrichment sequencing data was available (Yang et al., 2018), was re-analyzed in this study. Reported ploidy levels were variable, ranging from 2 to 12 according to the accession. The panel was genotyped using HaploCharmer for the same set of 86,681 genomic regions.

The same parameters as those used for the R570 progeny were used, with the exception of the minimum read depth “min_gt_depth” set to 10, due to constraints related to sequencing depth. Because the genomic regions analyzed to build the genetic map were, for a large part, defined based on the set of 60K probes defined by Yang et al. (2018) to analyse their diversity panel, most of the haplotypes could be recovered. A total of 1,854,032 haplotypes distributed in 60,429 phase sets were obtained from the raw presence-absence HPA output file.

#### Diversity analysis

Principal coordinate analyses (PCoAs) were performed on the diversity panel from the 180,115 haplotypes with less than 5% missing values coded as read depth ratios (i.e. haplotype depth to the total depth), and by excluding 6 accessions with more than 50% missing values. Remaining missing values were imputed using the average haplotype read depth ratios and PCoAs were calculated from an Euclidean distance matrix.

An ancestral origin was assigned to mapped markers by determining haplotypes private to *S. officinarum* or *S. spontaneum* accessions, i.e. at least one accession of the group genotyped as present for the haplotype but none from the other group. For *S. spontaneum*, a few accessions were discarded as they were likely introgressed with *S. robustum/S. officinarum* (see below).

## Results and discussion

### Analysis of the self-progeny of the polyploid cultivar R570

Haplocharmer was applied to whole genome sequencing data of a self-progeny of the R570 sugarcane cultivar (Grivet et al., 1994; Grivet et al., 1996; Healey et al., 2024) to evaluate its ability to accurately genotype haplotypes from 80K genomic regions in a segregating population. Among the 215,704 haplotypes obtained, the estimated proportion of false positives was below 1% (Table 1), demonstrating the high quality of the haplotyping obtained with HaploCharmer. A haplotype was qualified as false positive if its expected proportion in the progenies genotyped was low (*δ*_*k*_ = 2.25%), which is a very unlikely segregation for true haplotypes. These false positives likely resulted from sequencing and mapping errors. Similar proportions were found for SNPs and indels contributing to haplotypes.

**Table 1.** Percentage *π*_*k*_ of markers (haplotypes or alleles) from each category (false positive, single-dose or multi-dose) and expected percentage *δ*_*k*_ of progeny genotyped as present in the category, according to the type of marker: haplotypes obtained from HaploCharmer or each type of variant constituting the haplotypes (bi- and multi-allelic SNPs, bi-allelic indels and other more complex variants).

|  | False positive |  | Single-dose |  | Multi-dose |  |
| --- | --- | --- | --- | --- | --- | --- |
| | $\pi_k$ | $\delta_k$ | $\pi_k$ | $\delta_k$ | $\pi_k$ | $\delta_k$ |
| Haplotypes | 0.82% | 2.25% | 39.00% | 72.86% | 60.18% | 99.31% |
| Bi-allelic SNPs | 0.43% | 2.24% | 23.10% | 73.72% | 73.47% | 99.74% |
| Bi-allelic indels | 0.30% | 1.80% | 23.84% | 73.84% | 75.86% | 99.75% |
| Multi-allelic SNPs | 0.61% | 2.20% | 30.91% | 72.76% | 68.48% | 99.52% |
| Others | 0.73% | 2.84% | 30.05% | 72.83% | 69.22% | 99.39% |

The estimated proportion of single-dose haplotypes was 39.00%, which was higher than the proportion obtained with bi-allelic SNPs (23.10%), bi-allelic indels (23.84%), multi-allelic SNPs (30.91%) or other variants (30.05%). This result can be explained by multi-dose SNPs or indels which, when combined, can reveal single-dose haplotypes. While the theoretical proportion of single-dose markers present in a self-progeny is 75%, a lower proportion (*δ*_*k*_ *≈* 73%) was observed for haplotypes, SNPs and indels. This lower proportion could result from a slight segregation bias due to meiosis abnormalities or insufficient sequencing depth leading to erroneous genotyping of haplotypes or variants from which they were defined.

Multi-dose haplotypes were genotyped as present in nearly all progenies (*δ*_*k*_ = 99.31%), and the same was observed for SNPs and indels contributing to haplotypes. This category aggregated different dosage configurations (e.g. double-dose, triple-dose, …). It was not realistic to differentiate them due to (i) the limited size of our progeny sample and (ii) the conditioning of the expected proportion of multidose markers genotyped as present (*δ*_*k*_) on the pairing configuration at meiosis. This means that two double-dose haplotypes may not have the same *δ*_*k*_ value if they are located on chromosomes with contrasting pairing affinities at meiosis, although this value is close to 100% in both cases.

The 81,857 single-dose haplotypes could be grouped into 25,094 non-redundant bins used as effective markers for genetic mapping. In comparison, 21,789 and 24,435 non-redundant bins could be obtained by considering single-dose bi-allelic SNPs or all variants included in haplotypes, respectively. This higher number of effective markers obtained from haplotypes directly results from the phenomenon where single-dose haplotypes can emerge from variants whose alleles are all multi-dose.

Using haplotypes, 152 cosegregation groups of more than 20 haplotypes were obtained and the 18 corresponding to copies of chromosome 1 were represented in Figure 2. The physical position of haplotypes on the monoploid reference sequence was based on the coordinates of the PS from which they were identified. In addition, the consistency of these cosegregation groups was validated using haplotype pairwise recombination fractions (Figure S2). Although 12 chromosome-scale cosegregation groups are expected for R570 for chromosome 1 (Piperidis and D’Hont, 2020), the cosegregation groups obtained did not sum to 12 copies at each position along chromosome 1. This partial representation of the genome obtained from the linkage map was expected since large chromosome segments are identical by descent due to inbreeding in the genealogy of cultivar R570 (Healey et al., 2024), resulting in regions of the genome where haplotypes are multi-dose (i.e. no single dose were available for genetic mapping).

**Figure 2.**
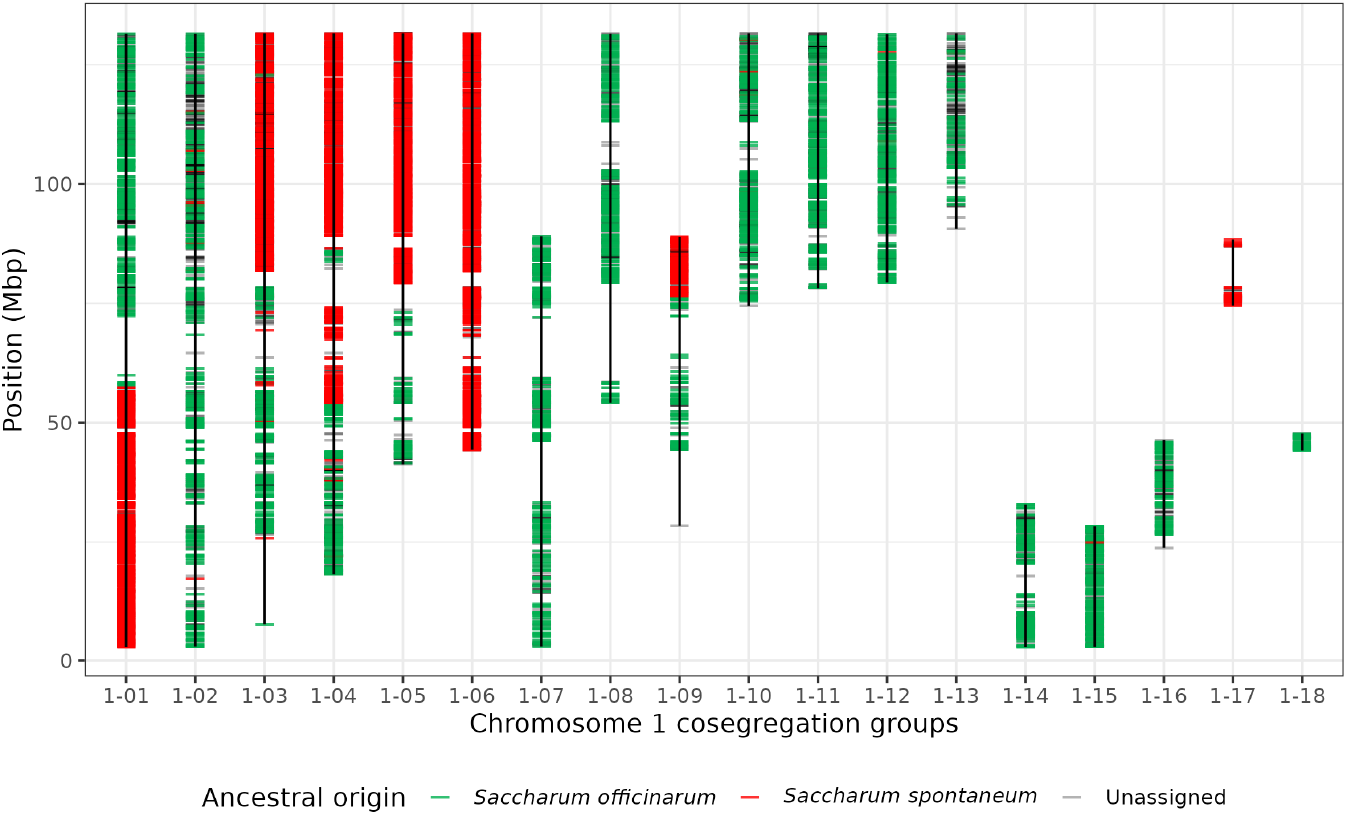
Cosegregation groups obtained from single-dose haplotypes for the chromosome 1 homology class. Each haplotype is represented by a segment colored according to its ancestral origin. Only cosegregation groups of at least 20 haplotypes are displayed.

Modern sugarcane cultivars such as R570 resulted from hybridization between two species with distinct basic chromosome numbers, *S. officinarum* (x=10) and *S. spontaneum* (x=8). In *S. spontaneum*, two sets of three chromosomes have each been rearranged into two chromosomes, in comparison with the ancestral chromosome structure observed in *S. officinarum* (Garsmeur et al., 2018; Piperidis and D’Hont, 2020). Accordingly, some cosegregation groups corresponded to known rearranged chromosomes. To illustrate, two cosegregation groups involving chromosomes 5, 6 and 9, which have been rearranged into two in *S. spontaneum*, are presented in Figure 3. The two cosegregation groups and the positioning of haplotypes along the monoploid reference assembly (ancestral genome structure) on Figure 3 well reflect this known rearrangement, with one part of chromosome 9 merged with chromosome 5 and the other part merged with chromosome 6.

**Figure 3.**
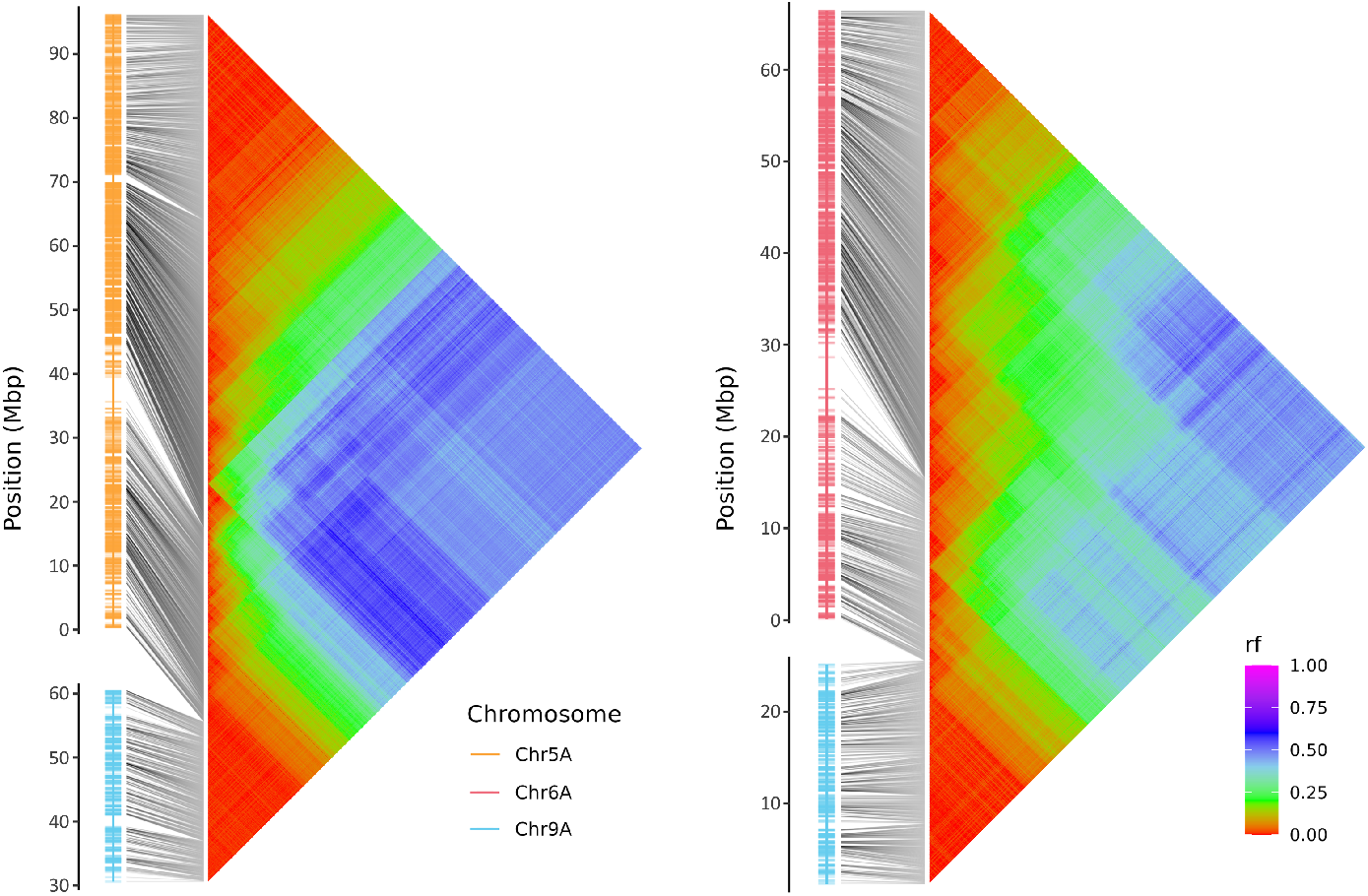
Two cosegregation groups corresponding to *S. spontaneum* rearranged chro-mosomes 5/9 and 6/9, with haplotypes positioned on the R570 reference (Chr5A, Chr6A and Chr9A) along with heatmaps of pairwise recombination fractions (rf) estimated between haplotypes. The positioning of haplotypes along the reference corresponds to the ancestral genome structure.

### Analysis of the polyploid *Saccharum* diversity panel

The diversity panel of 307 *Saccharum* and related genera accessions presented by Yang et al. (2018) was characterized for the same PSs. An haplotype-based PCoA including all accessions clearly distinguished non-*Saccharum* accessions from *Saccharum* accessions (Figure 4A). Within *Saccharum* accessions, a second PCoA opposed *S. robustum* and *S. officinarum* to *S. spontaneum* on the first axis, while the second axis mainly reflected *S. spontaneum* variation (Figure 4B). A few *S. spontaneum* and *S. robustum* accessions appeared shifted toward the center of the PCoA, suggesting that they are admixed, as already reported in Yang et al. (2018). As expected, declared hybrids including *S. barberi* and *S. sinense* were positioned between their parental species on the PCoAs (i.e. *S. robustum*/*S. officinarum* and *S. spontaneum*).

**Figure 4.**
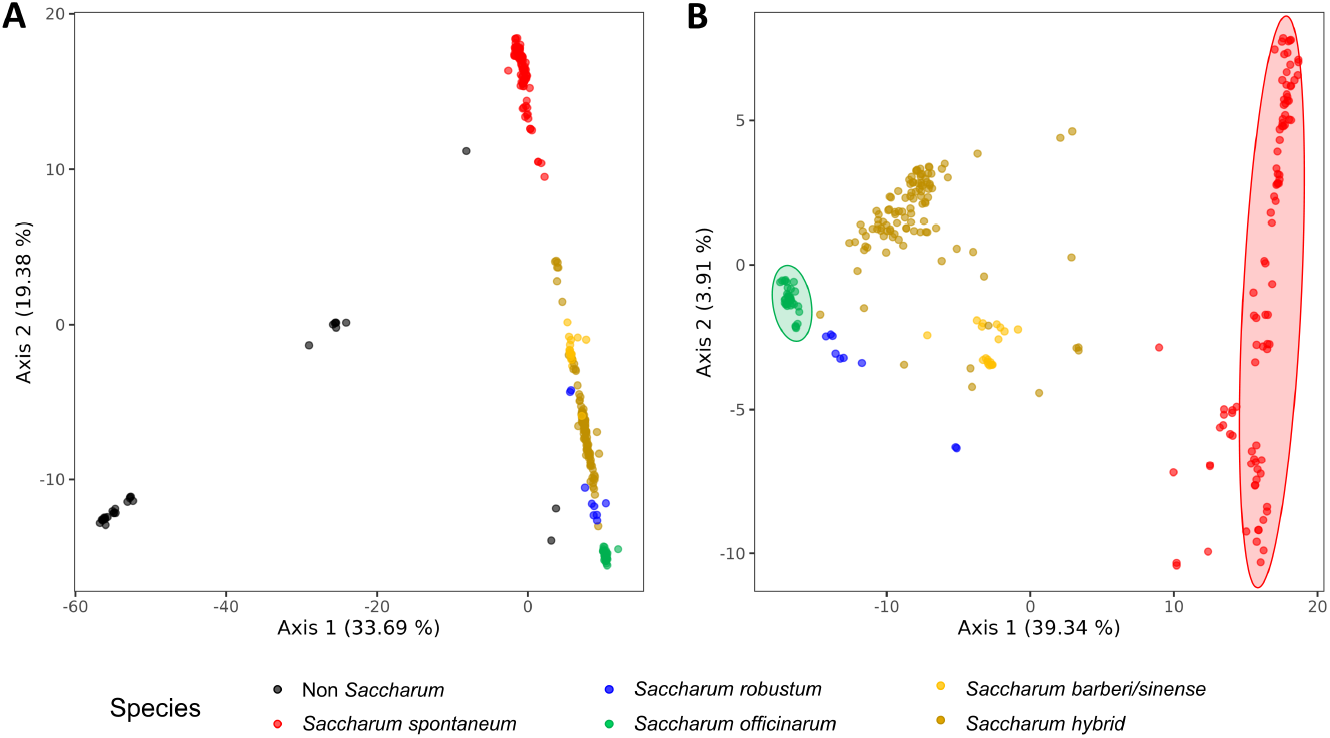
Principal coordinate analyses of (A) all accessions from the diversity panel of Yang et al. (2019) and (B) only *Saccharum* accessions. Dots are colored according to the *Saccharum* species. The colored ellipses highlight the set of *S. officinarum* and *S. spontaneum* accessions used to infer an ancestral origin to haplotypes of the genetic map.

Ancestral origins of the single-dose haplotypes used to build the genetic map could be assigned by determining those private to pure *S. officinarum* or *S. spontaneum* accessions. They revealed clear ancestry blocks in the genetic map of chromosome 1 (Figure 2), in accordance with the existence of recombinant chromosomes between the two parental species in sugarcane modern cultivars (Piperidis and D’Hont, 2020).

### Implementation

An important aspect of the workflow concerns the definition of preconfigured genomic regions (i.e. PSs). For whole-genome sequencing data, they can be randomly defined in non-repetitive sequences or in sequences of putative interest (e.g. exons or cis-regulatory elements). For genotyping-by-sequencing approaches, PSs can be defined in read-enriched regions. Alternatively, an automatic approach can be envisaged, consisting of targeting all adjacent genomic regions of a given size across the entire genome, after masking repetitive elements.

In addition, the choice of PS size is critical as it has an impact on the number of sequencing reads that are retained to define haplotypes. Indeed, only sequencing reads that cover the PSs are considered, resulting in a negative relationship between PS size and read depth of genotypic datapoints (Figure S3). To accurately genotype haplotypes, PS size should be chosen relative to sequencing depth and a minimum read depth should be fixed that prevents missing low frequency haplotypes when the read depth is too low, e.g. >10% of single-dose haplotypes missed when DP<30 in the 12x R570 sugarcane cultivar (Figure S4).

## Conclusions

This study highlights the effectiveness of HaploCharmer in accurately haplotyping populations of varying complexity in terms of structure (segregating population or diversity panel) or ploidy. HaploCharmer thus offers a robust haplotype calling framework for diversity, genetic mapping, and quantitative genetics studies in both diploid and polyploid species.

## Acknowledgments

A preprint version of this article has been peer-reviewed and recommended by Peer Community In Genomics (https://doi.org/10.24072/pci.genomics.100434; Campos Dominguez, 2025).

## Fundings

This work was supported by the International Consortium for Sugarcane Biotechnology (Project ICSB37). Whole genome sequencing data of the R570 self-progeny were produced by the Joint Genome Institute (https://ror.org/04xm1d337, proposal: https://doi.org/10.46936/10.25585/60001084), a USA Department of Energy (DOE) Office of Science User Facility and the DOE Joint BioEnergy Institute, which are supported by the Office of Science of the DOE operated under Contract No. DE-AC02-05CH11231.

## Conflict of interest disclosure

The authors declare that they have no conflicts of interest in relation to the content of the article.

## Data, script, code, and supplementary information availability

HaploCharmer source code is available at https://gitlab.cirad.fr/agap/seg/haplocharmer, along with documentation and examples. It can easily be installed on any Linux system. Dependencies are managed automatically via a conda installation or using a docker image.

Whole-genome sequencing data of the R570 progeny is available from the Joint Genome Institute portal: https://genome.jgi.doe.gov/portal/Undpolgesequence/Undpolgesequence.info.html. The target-enrichment sequencing reads of the diversity panel are available from the NCBI Short Reads Archive (project number: SRP132365). The genome assembly of R570 was available from the Phytozome portal: https://phytozome-next.jgi.doe.gov/info/Sofficinarumxspontanev2_1.

## Supplemental information

Figure S1: Number of haplotypes called (y-axis) in each progeny (x-axis). Colors correspond to distinct categories describing the presence, absence or missing information (NA) in comparison with R570, with R570 on the left, and the progeny on the right. For instance, “presence - absence” (in orange) corresponds to a haplotype present in R570 and absent in the progeny.

Figure S2: Cosegregation groups corresponding to chromosome 1, with haplotypes positioned on the monoploid reference (Chr1A) along with heatmaps of pairwise recombination fractions (rf) estimated between haplotypes. The positioning of haplotypes along the reference corresponds to the ancestral genome structure.

Figure S3: Average read depth of R570 and its self-progeny over the 86,681 PSs with varying size: 1, 10, 30, 50 or 80 bp. For each PS, only sequencing reads that fully span the entire region are considered.

Figure S4: Percentage of missed single-dose haplotypes (simplex) according to read depth of the ge
notypic datapoint in R570. Read depths categories superior to 80 with less than 200 haplotypes were not displayed.

## Notes

https://genome.jgi.doe.gov/portal/Undpolgesequence/Undpolgesequence.info.html

https://www.ncbi.nlm.nih.gov/sra?term=SRP132365

### Competing Interest Statement

The authors have declared no competing interest.

### Summary of Updates

Line numbers were removed. A PCI Genomic badge has been added. Few typos have been corrected.

