## Supplementary material for "HaploCharmer: a Snakemake workflow for read-scale haplotype calling adapted to polyploids": Figure S1

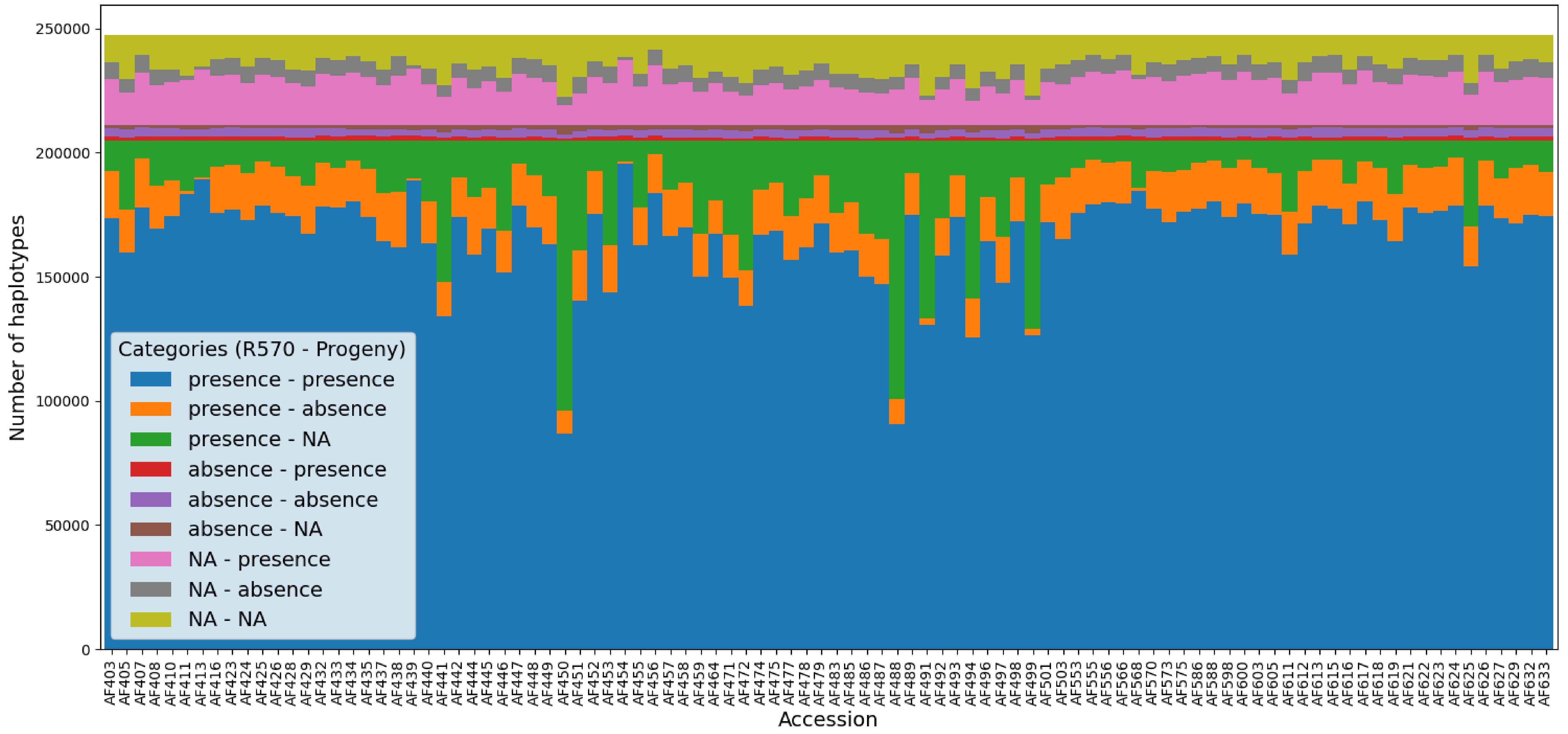


**Figure S1**: Number of haplotypes called (y-axis) in each progeny (x-axis). Colors correspond to distinct categories describing the presence, absence or missing information (NA) in comparison with R570, with R570 on the left, and the progeny on the right. For instance, “presence - absence” (in orange) corresponds to a haplotype present in R570 and absent in the progeny.
