## Supplementary material for "HaploCharmer: a Snakemake workflow for read-scale haplotype calling adapted to polyploids": Figure S2

# 1-01

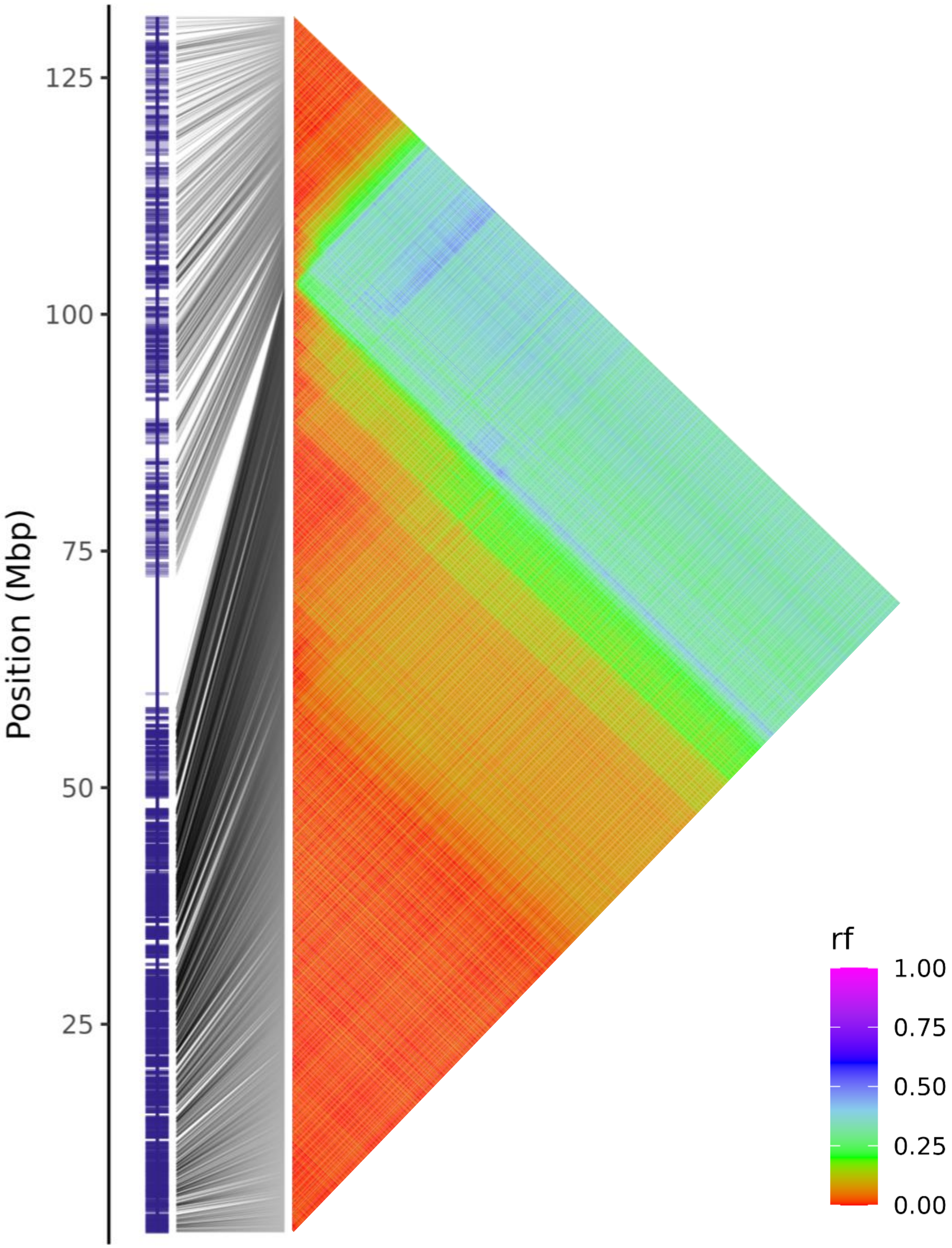


**Figure S2A**: Linkage group 1-01 of chromosome 1, with haplotypes positioned on the monoploid reference (Chr1A) along with heatmaps of pairwise recombination fractions (rf) estimated between haplotypes.

# 1-02

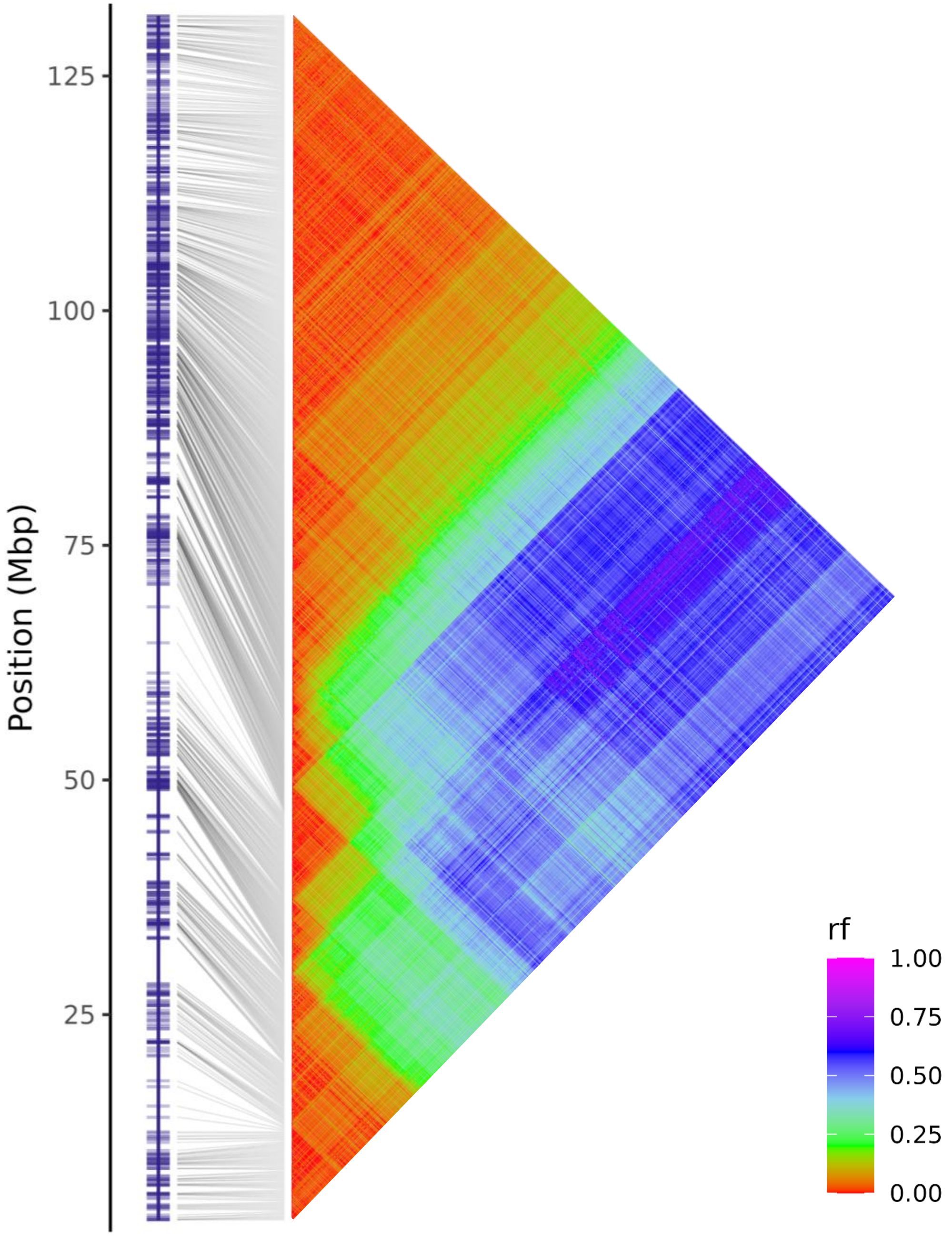


**Figure S2B**: Linkage group 1-02 of chromosome 1, with haplotypes positioned on the monoploid reference (Chr1A) along with heatmaps of pairwise recombination fractions (rf) estimated between haplotypes.

# 1-03

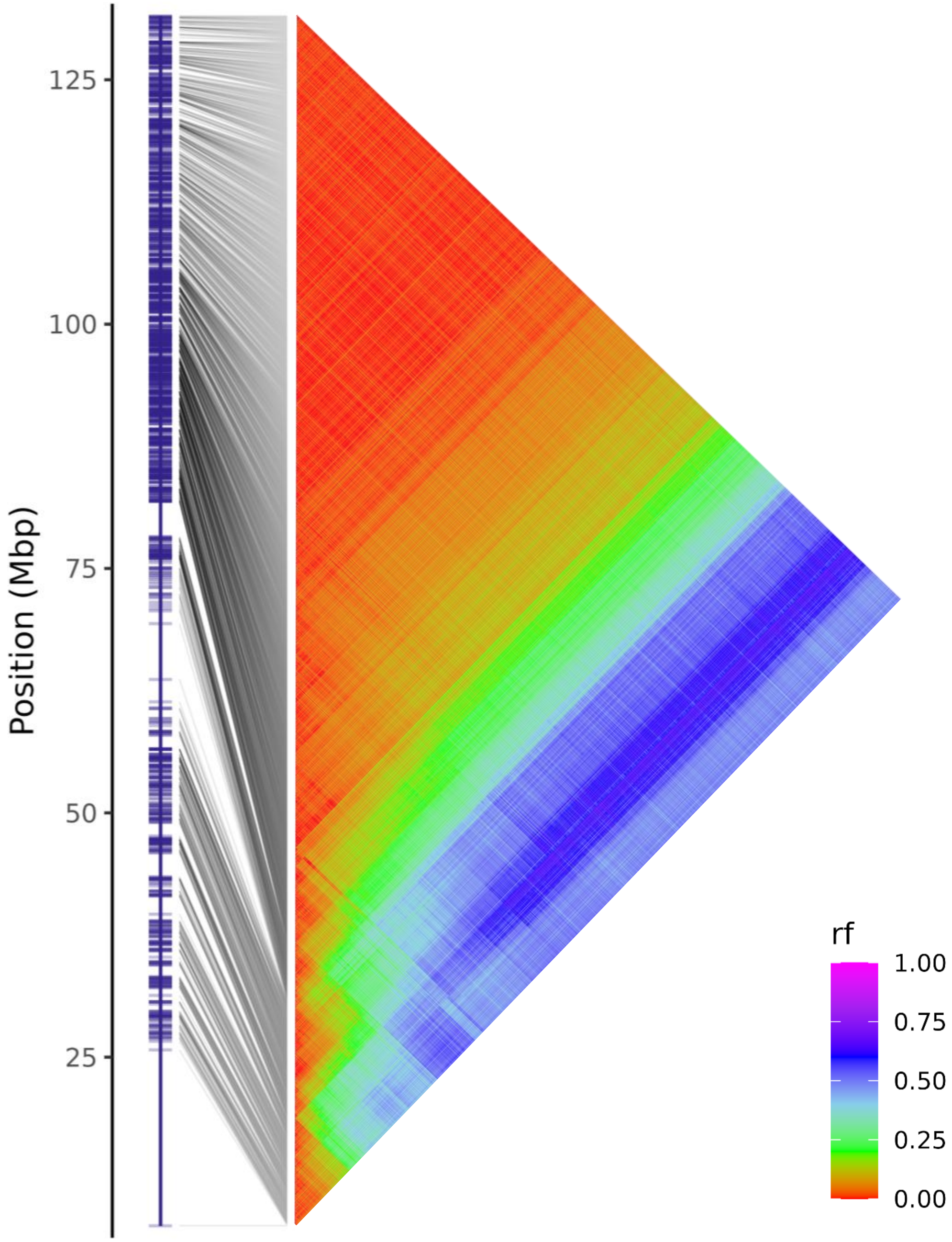


**Figure S2C**: Linkage group 1-03 of chromosome 1, with haplotypes positioned on the monoploid reference (Chr1A) along with heatmaps of pairwise recombination fractions (rf) estimated between haplotypes.

# 1-04

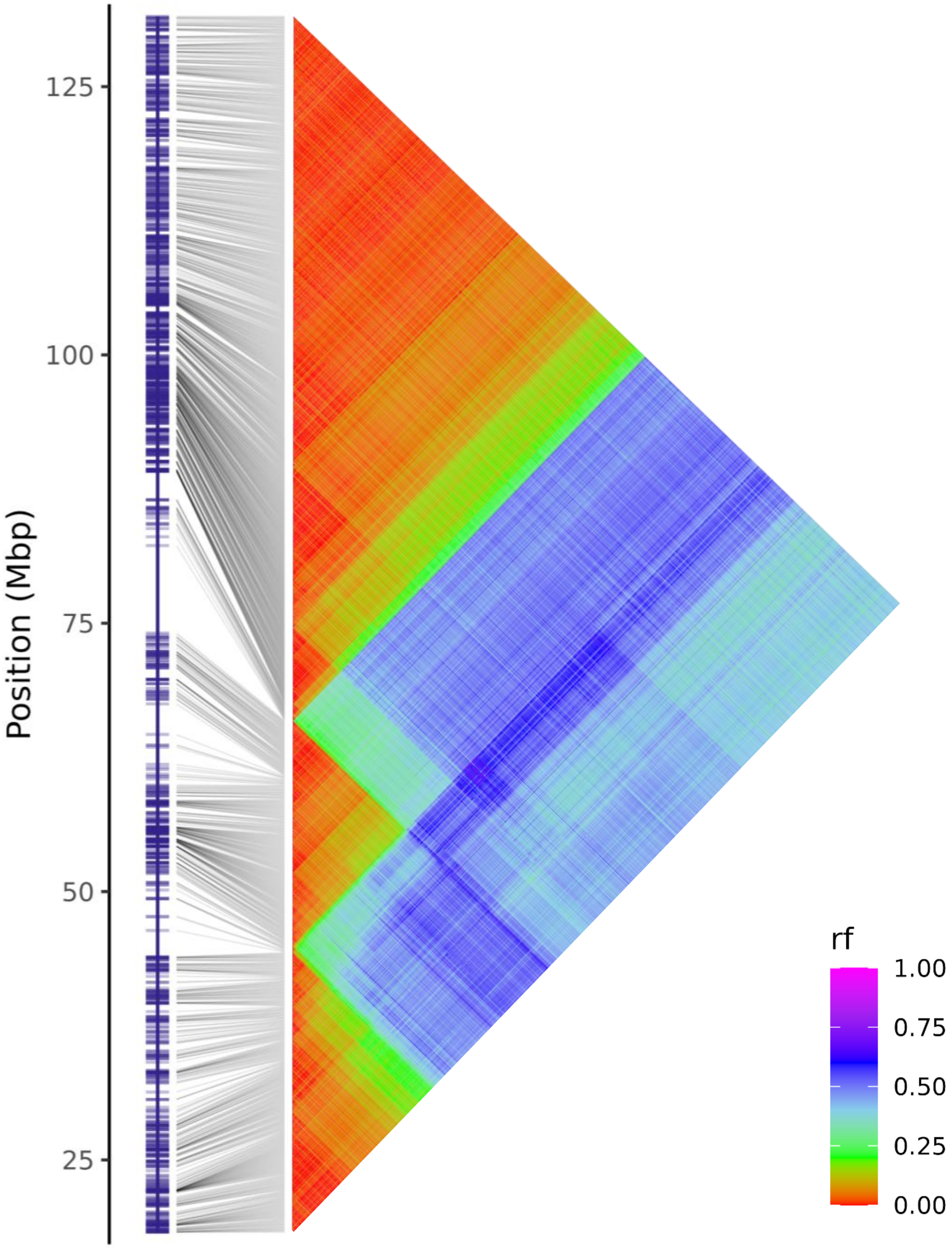


**Figure S2D**: Linkage group 1-04 of chromosome 1, with haplotypes positioned on the monoploid reference (Chr1A) along with heatmaps of pairwise recombination fractions (rf) estimated between haplotypes.

# 1-05

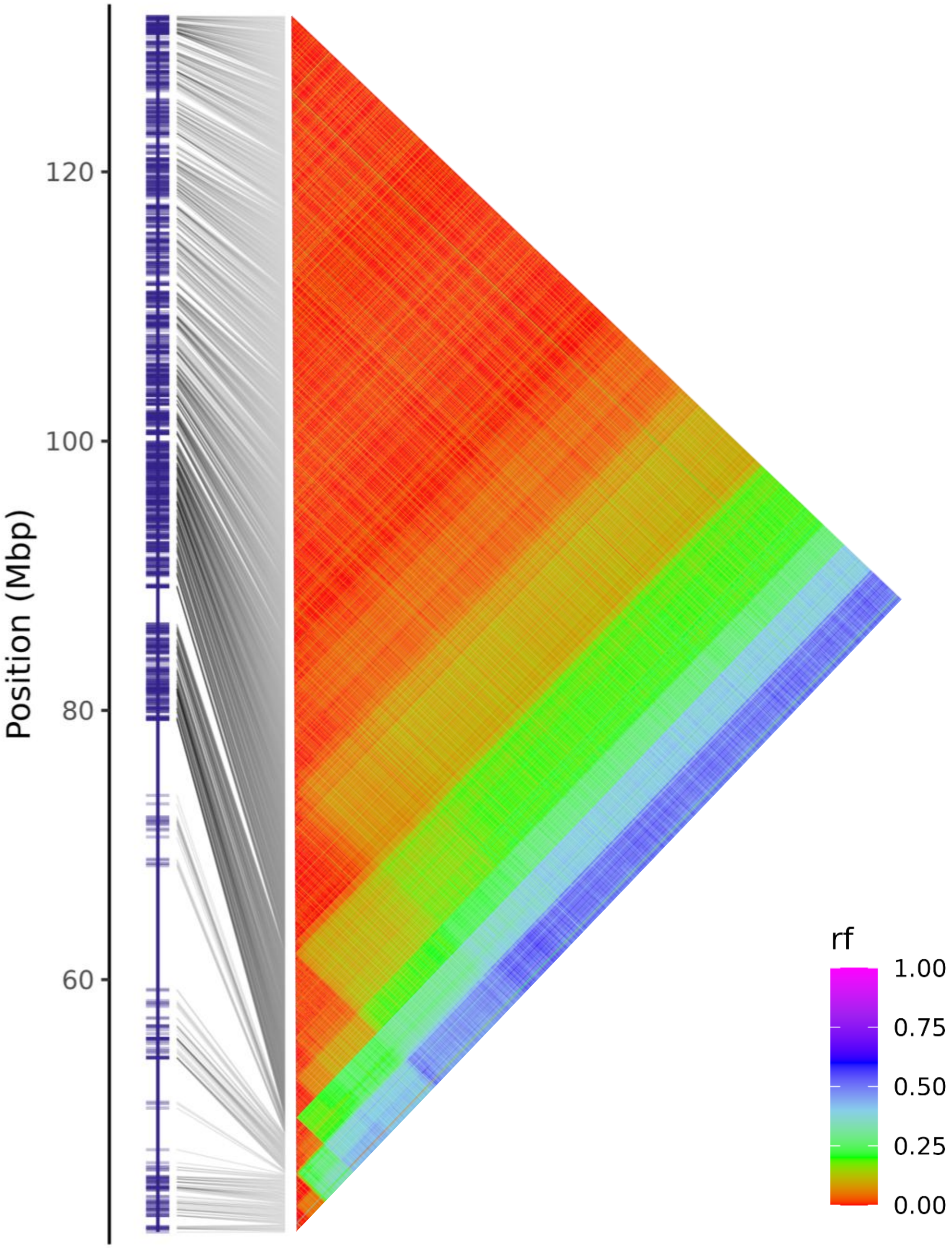


**Figure S2E**: Linkage group 1-05 of chromosome 1, with haplotypes positioned on the monoploid reference (Chr1A) along with heatmaps of pairwise recombination fractions (rf) estimated between haplotypes.

# 1-06

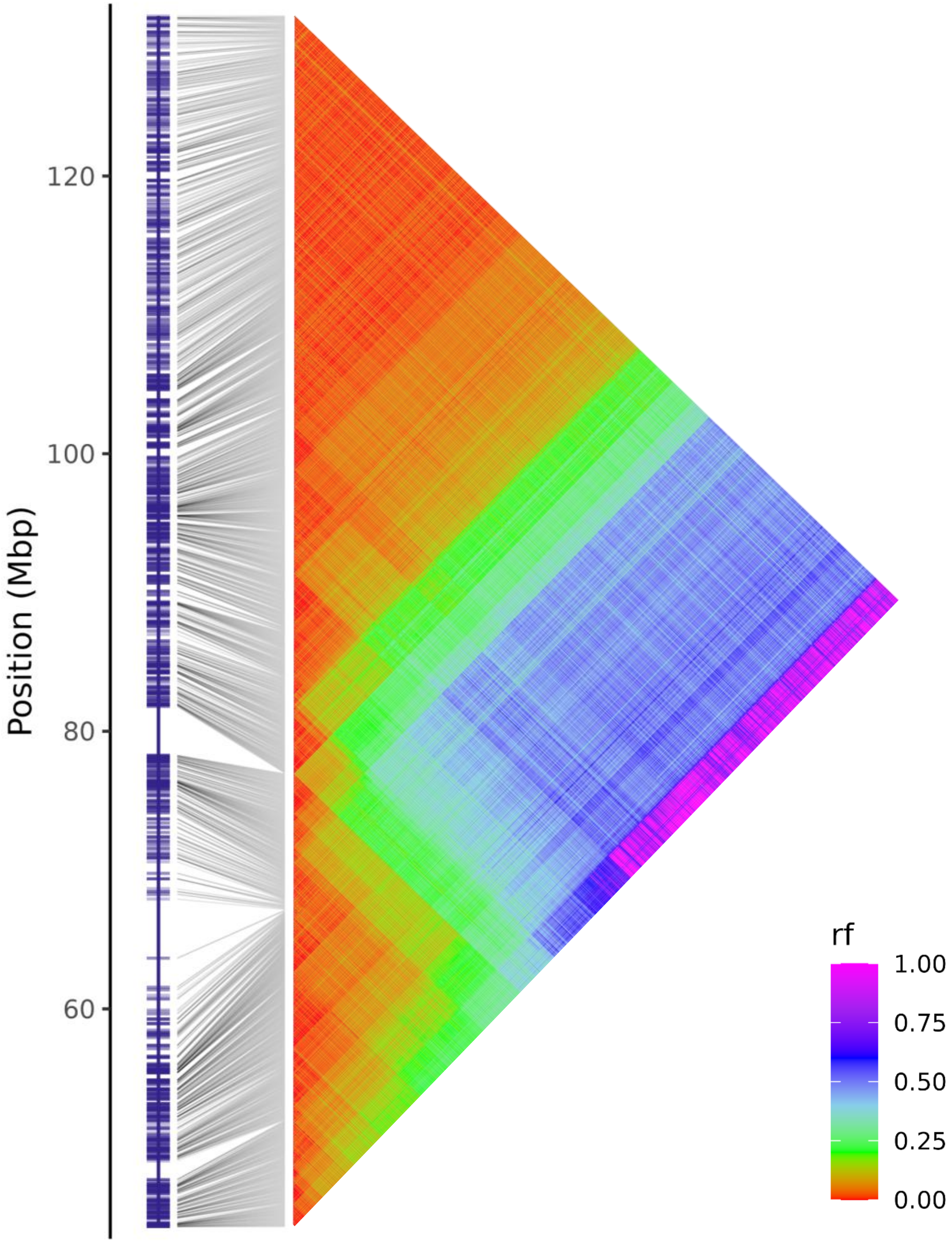


**Figure S2F**: Linkage group 1-06 of chromosome 1, with haplotypes positioned on the monoploid reference (Chr1A) along with heatmaps of pairwise recombination fractions (rf) estimated between haplotypes.

# 1-07

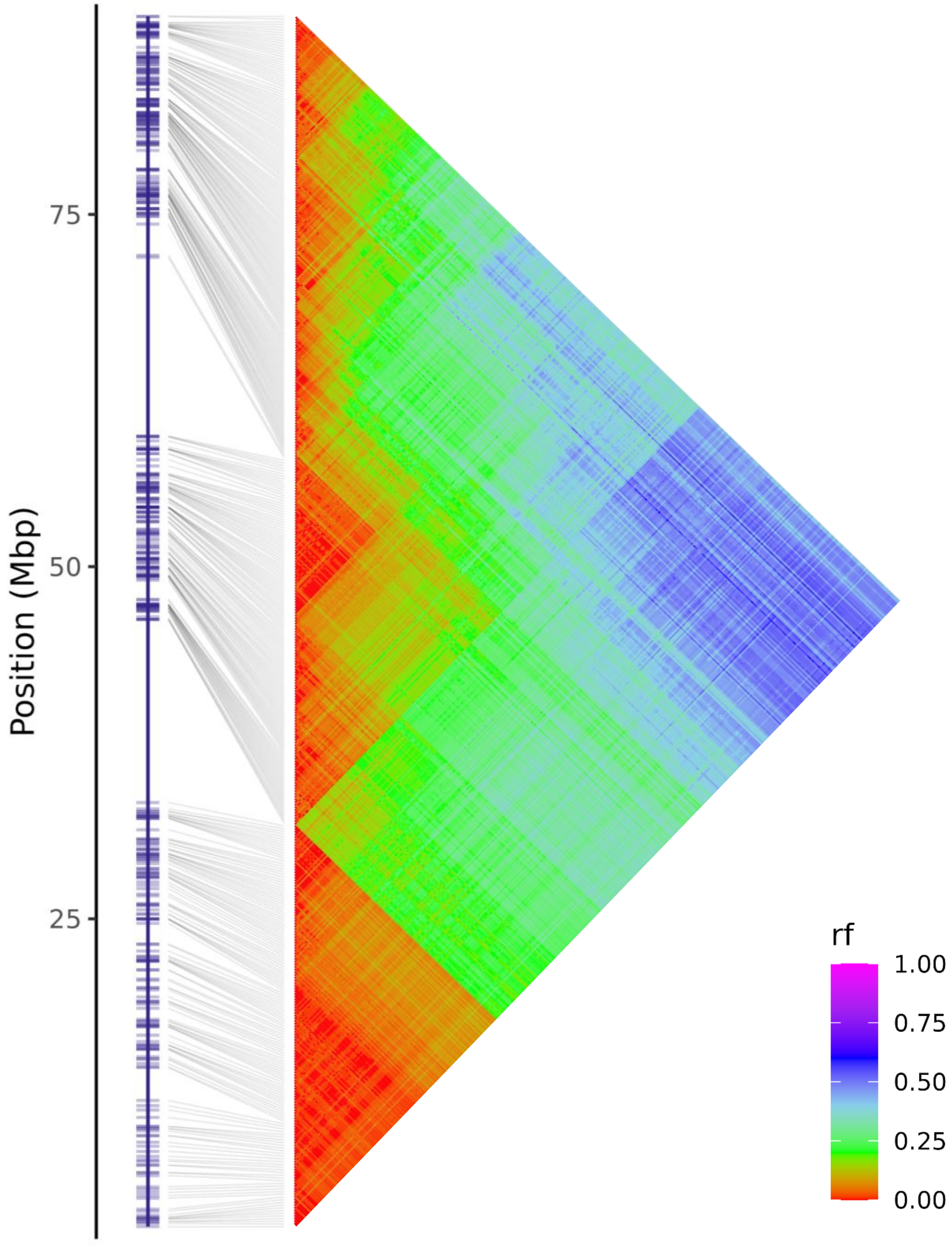


**Figure S2G**: Linkage group 1-07 of chromosome 1, with haplotypes positioned on the monoploid reference (Chr1A) along with heatmaps of pairwise recombination fractions (rf) estimated between haplotypes.

# 1-08

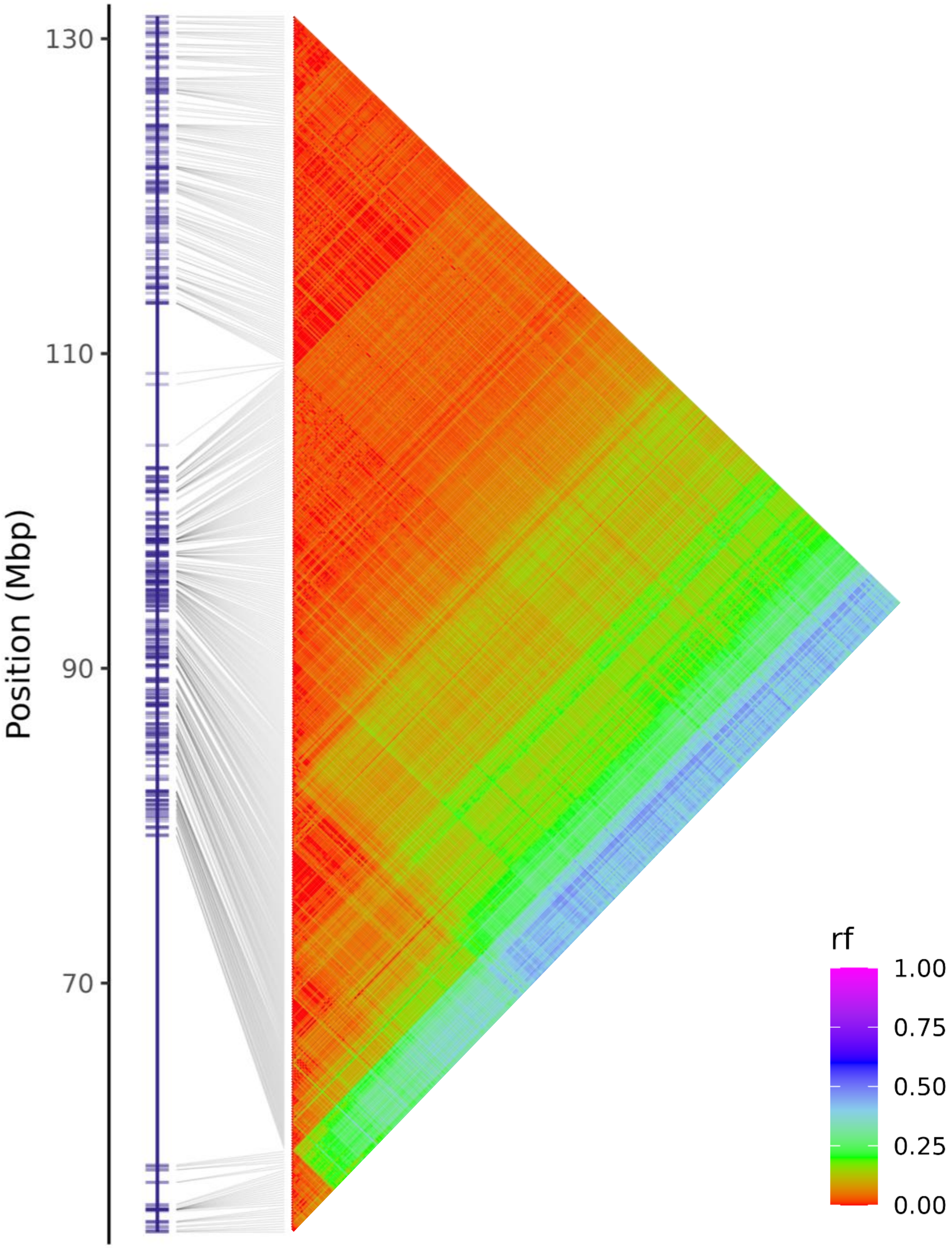


**Figure S2H**: Linkage group 1-08 of chromosome 1, with haplotypes positioned on the monoploid reference (Chr1A) along with heatmaps of pairwise recombination fractions (rf) estimated between haplotypes.

1-09

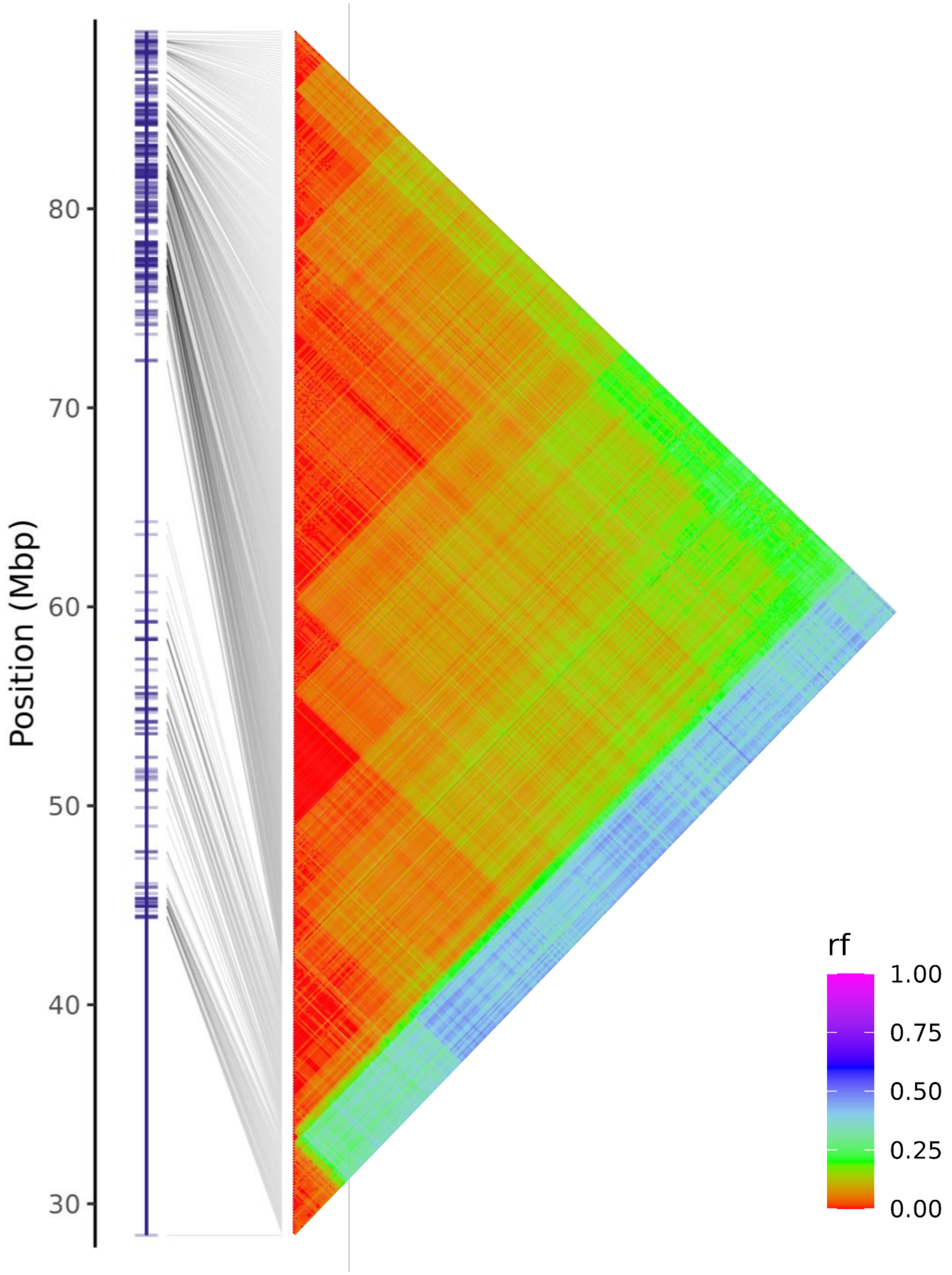


**Figure S2I**: Linkage group 1-09 of chromosome 1, with haplotypes positioned on the monoploid reference (Chr1A) along with heatmaps of pairwise recombination fractions (rf) estimated between haplotypes.

# 1-10

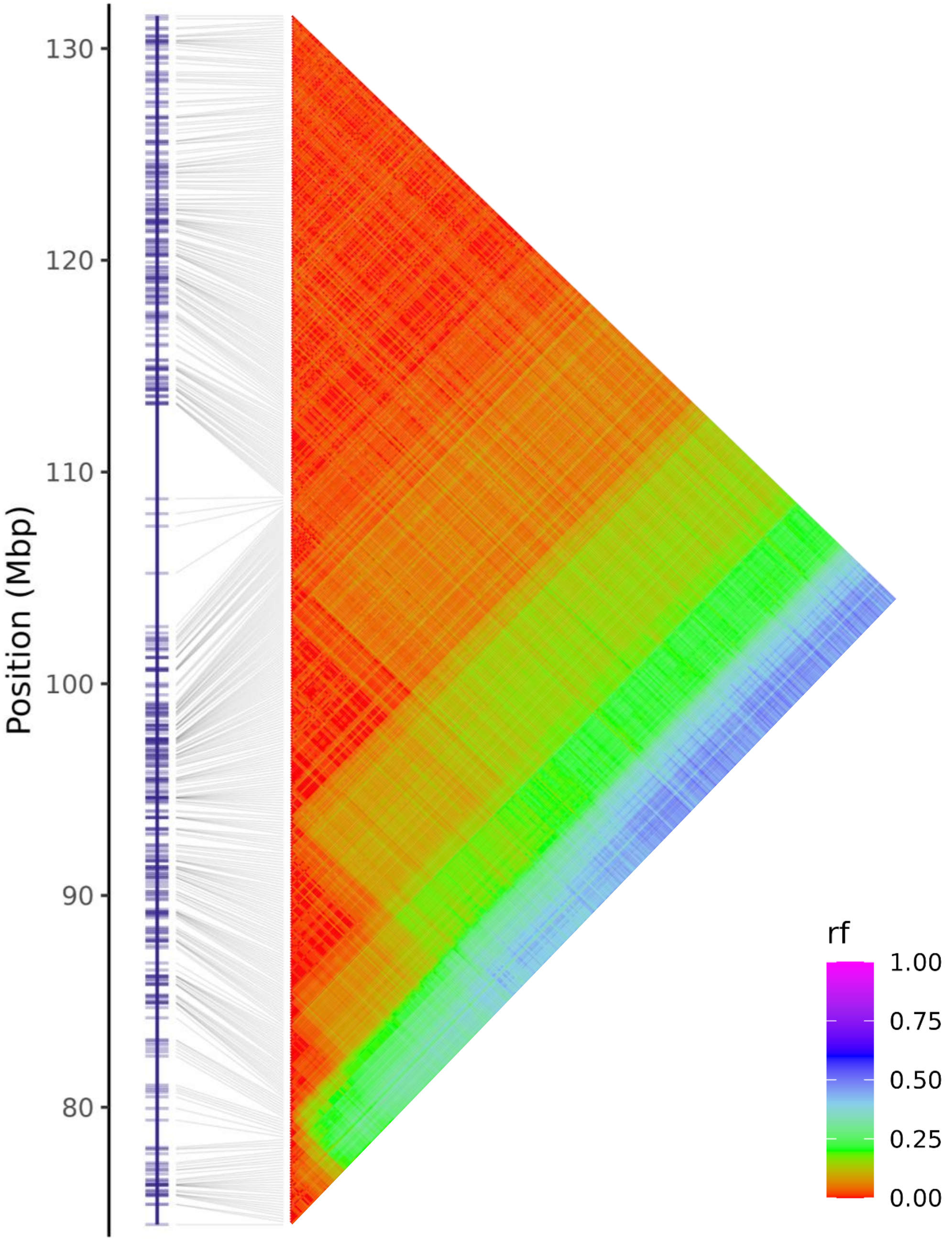


**Figure S2J**: Linkage group 1-10 of chromosome 1, with haplotypes positioned on the monoploid reference (Chr1A) along with heatmaps of pairwise recombination fractions (rf) estimated between haplotypes.

# 1-11

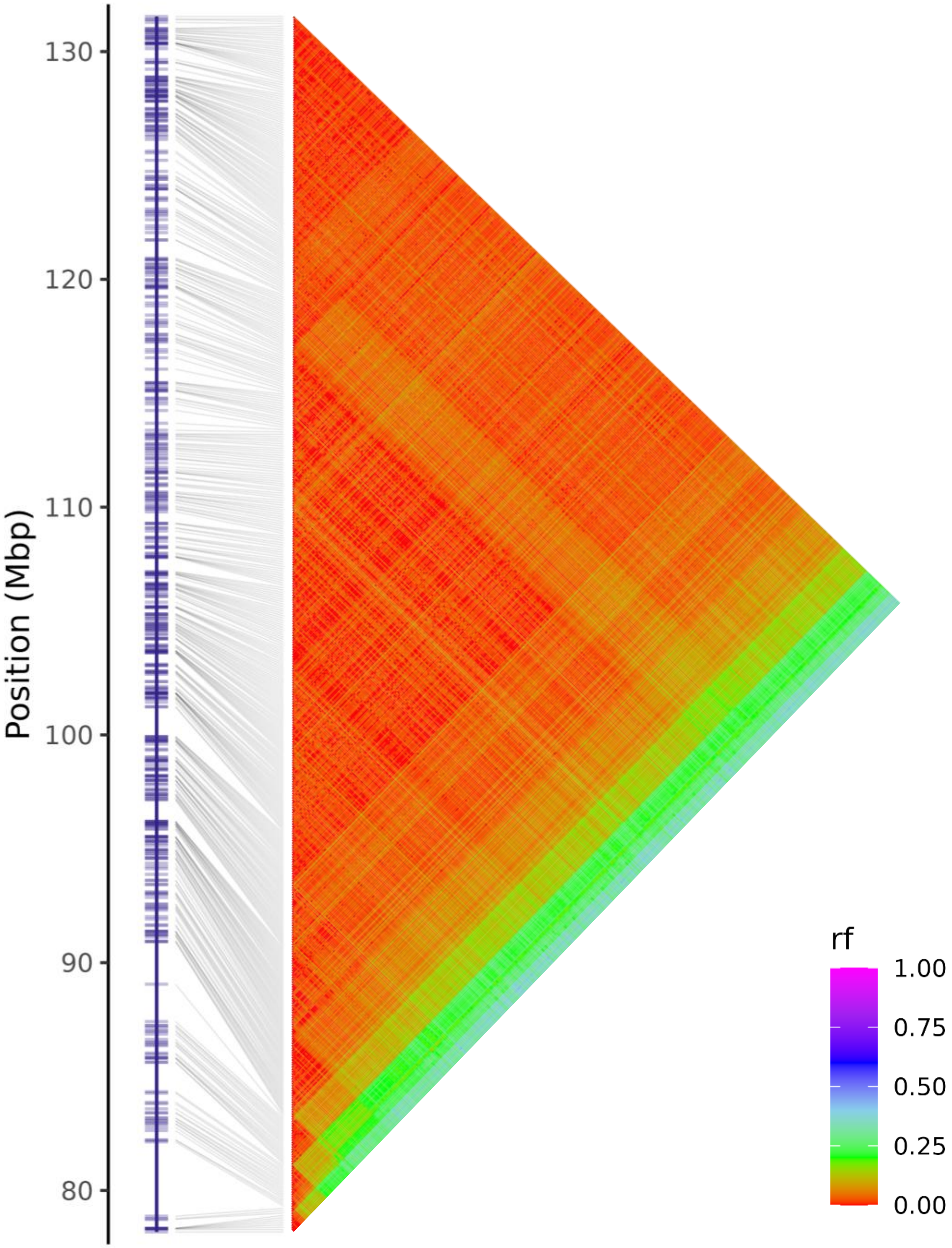


**Figure S2K**: Linkage group 1-11 of chromosome 1, with haplotypes positioned on the monoploid reference (Chr1A) along with heatmaps of pairwise recombination fractions (rf) estimated between haplotypes.

# 1-12

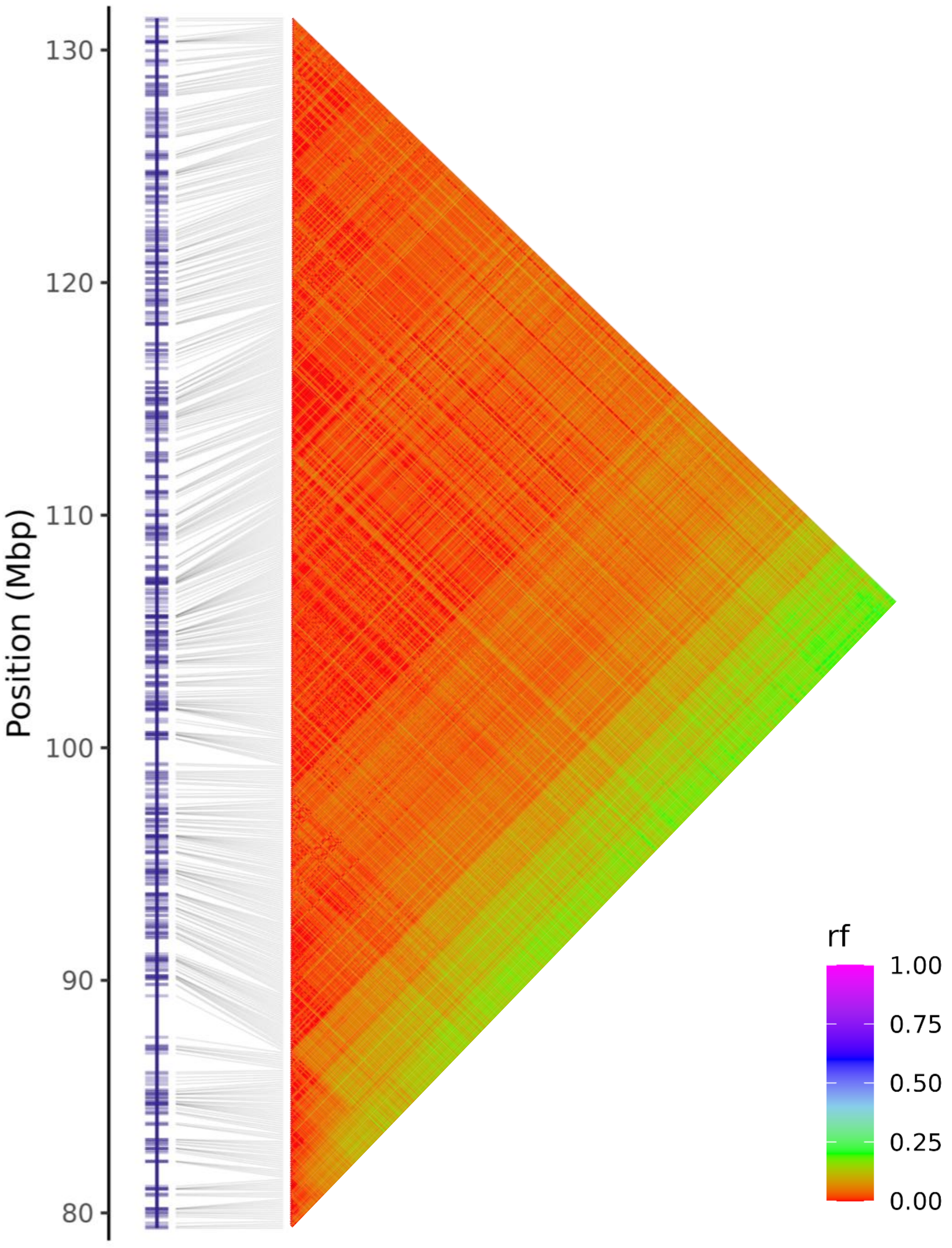


**Figure S2L**: Linkage group 1-12 of chromosome 1, with haplotypes positioned on the monoploid reference (Chr1A) along with heatmaps of pairwise recombination fractions (rf) estimated between haplotypes.

# 1-13

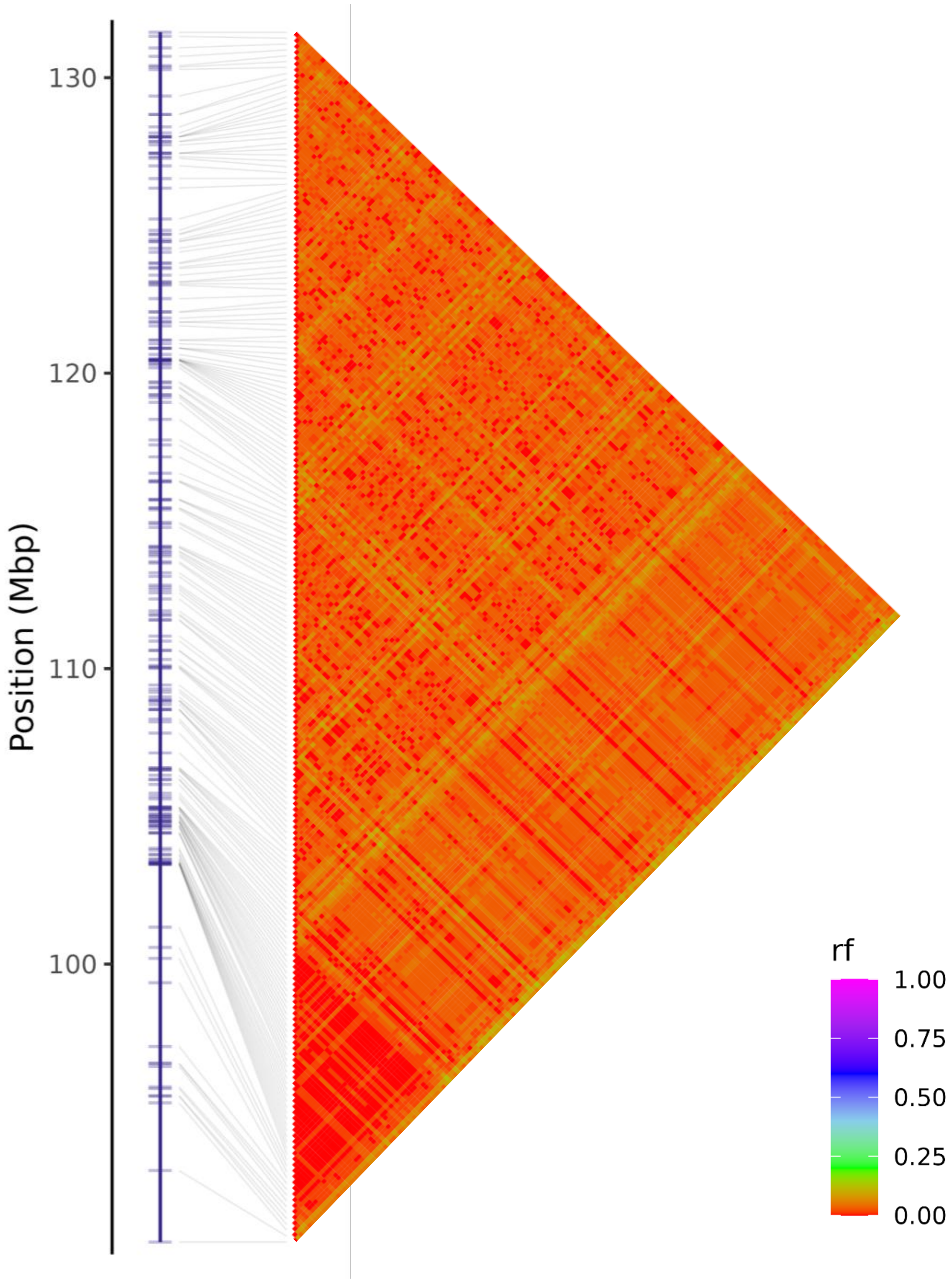


**Figure S2M**: Linkage group 1-13 of chromosome 1, with haplotypes positioned on the monoploid reference (Chr1A) along with heatmaps of pairwise recombination fractions (rf) estimated between haplotypes.

# 1-14

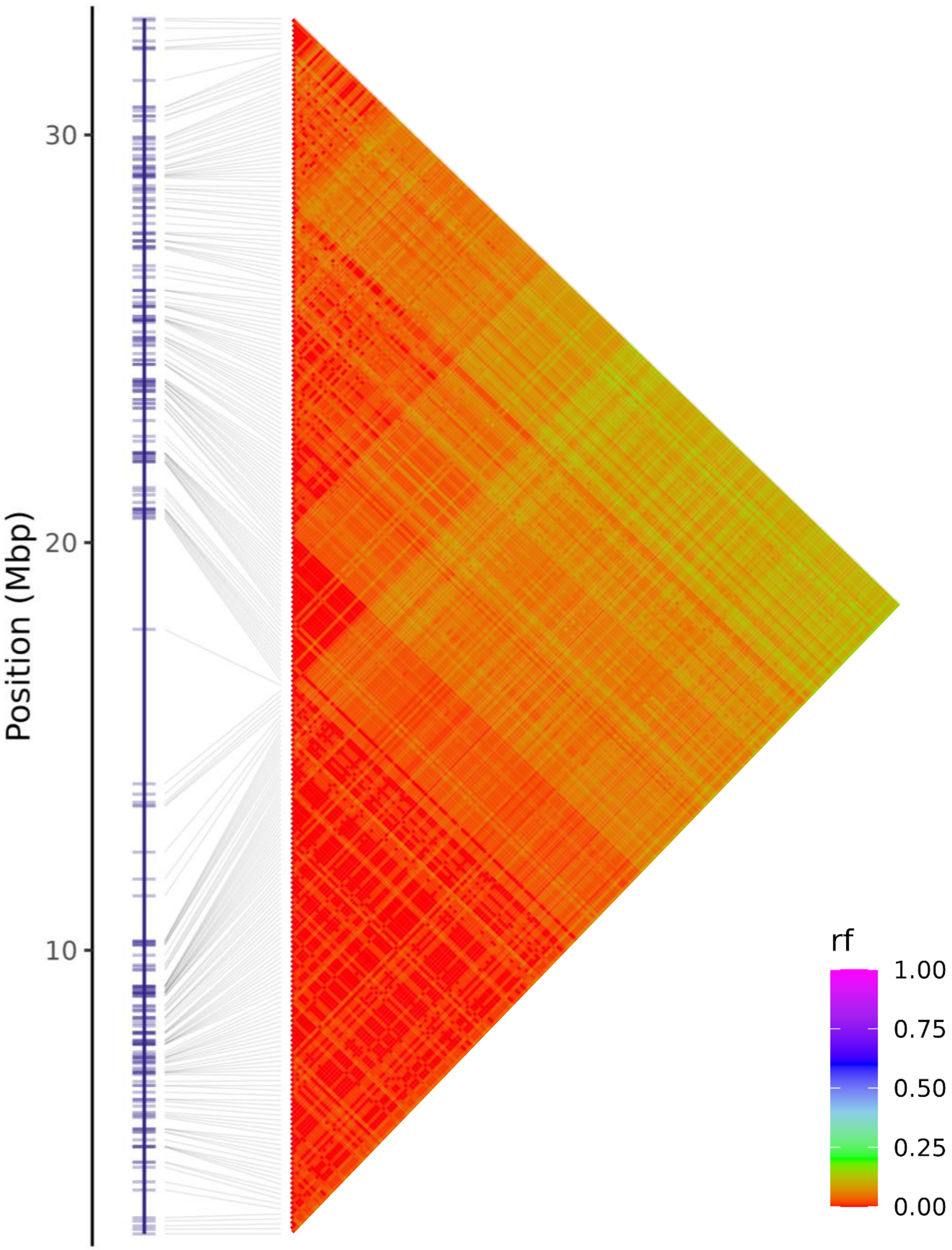


**Figure S2N**: Linkage group 1-14 of chromosome 1, with haplotypes positioned on the monoploid reference (Chr1A) along with heatmaps of pairwise recombination fractions (rf) estimated between haplotypes.

# 1-15

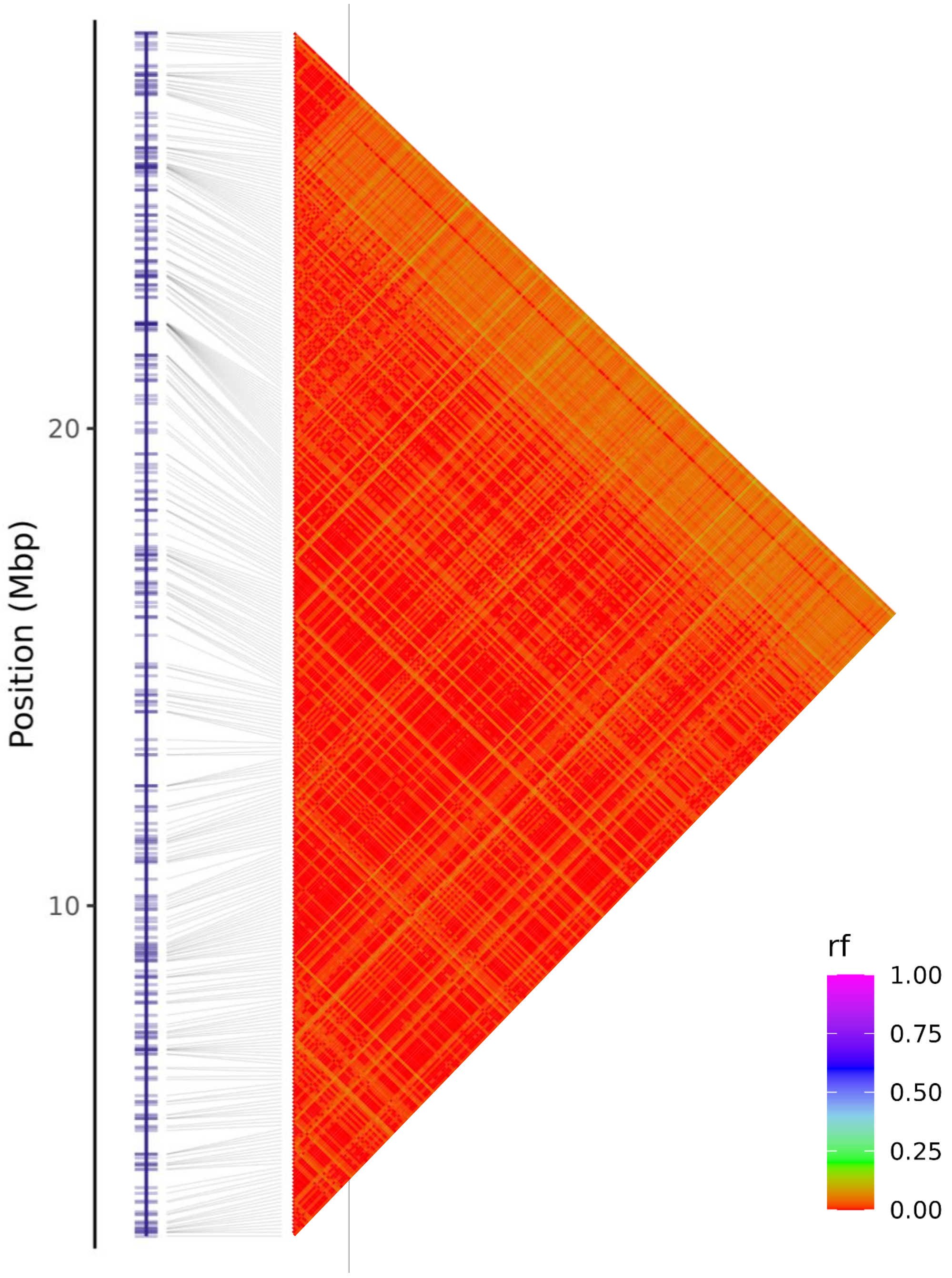


**Figure S2O**: Linkage group 1-15 of chromosome 1, with haplotypes positioned on the monoploid reference (Chr1A) along with heatmaps of pairwise recombination fractions (rf) estimated between haplotypes.

# 1-16

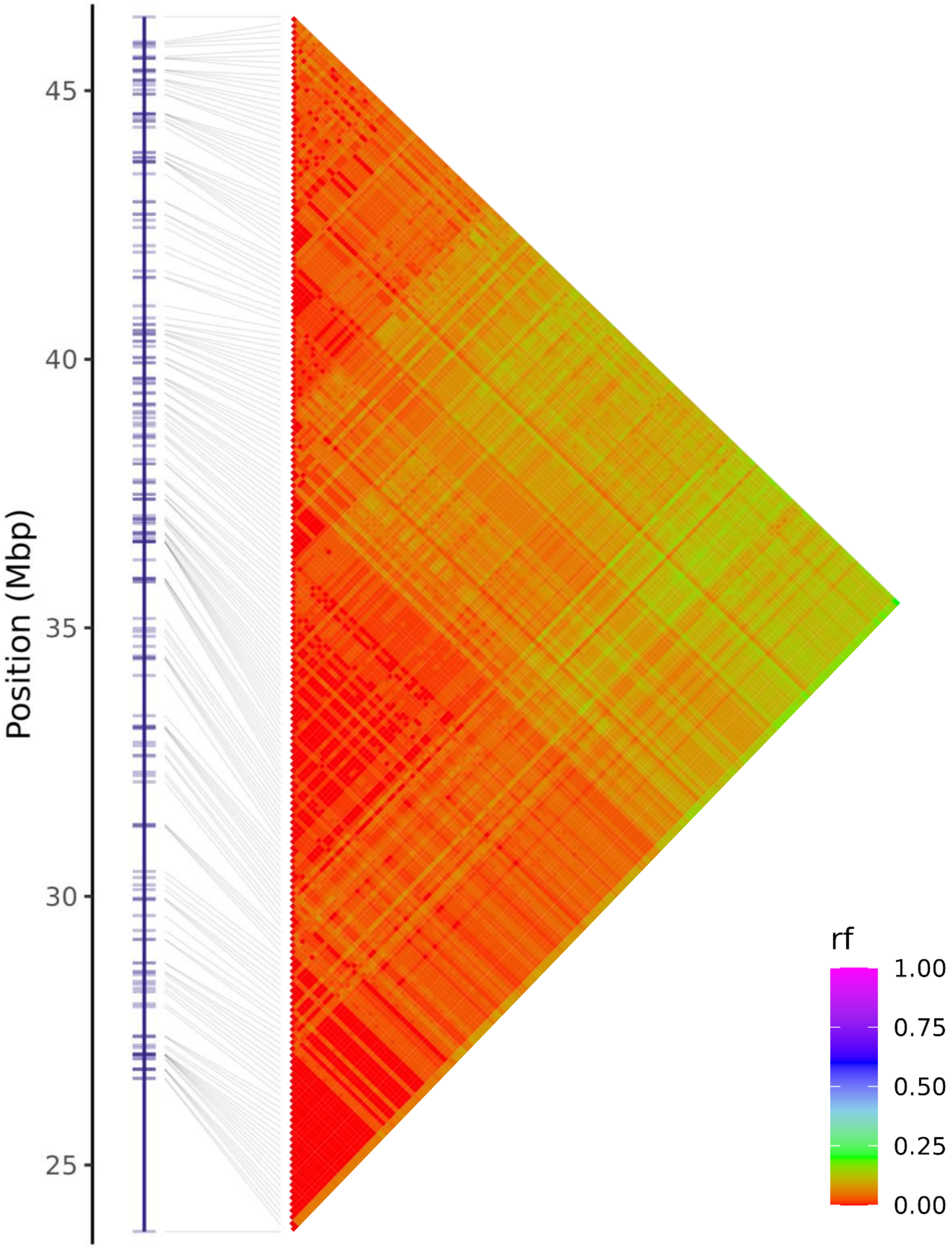


**Figure S2P**: Linkage group 1-16 of chromosome 1, with haplotypes positioned on the monoploid reference (Chr1A) along with heatmaps of pairwise recombination fractions (rf) estimated between haplotypes.

# 1-17

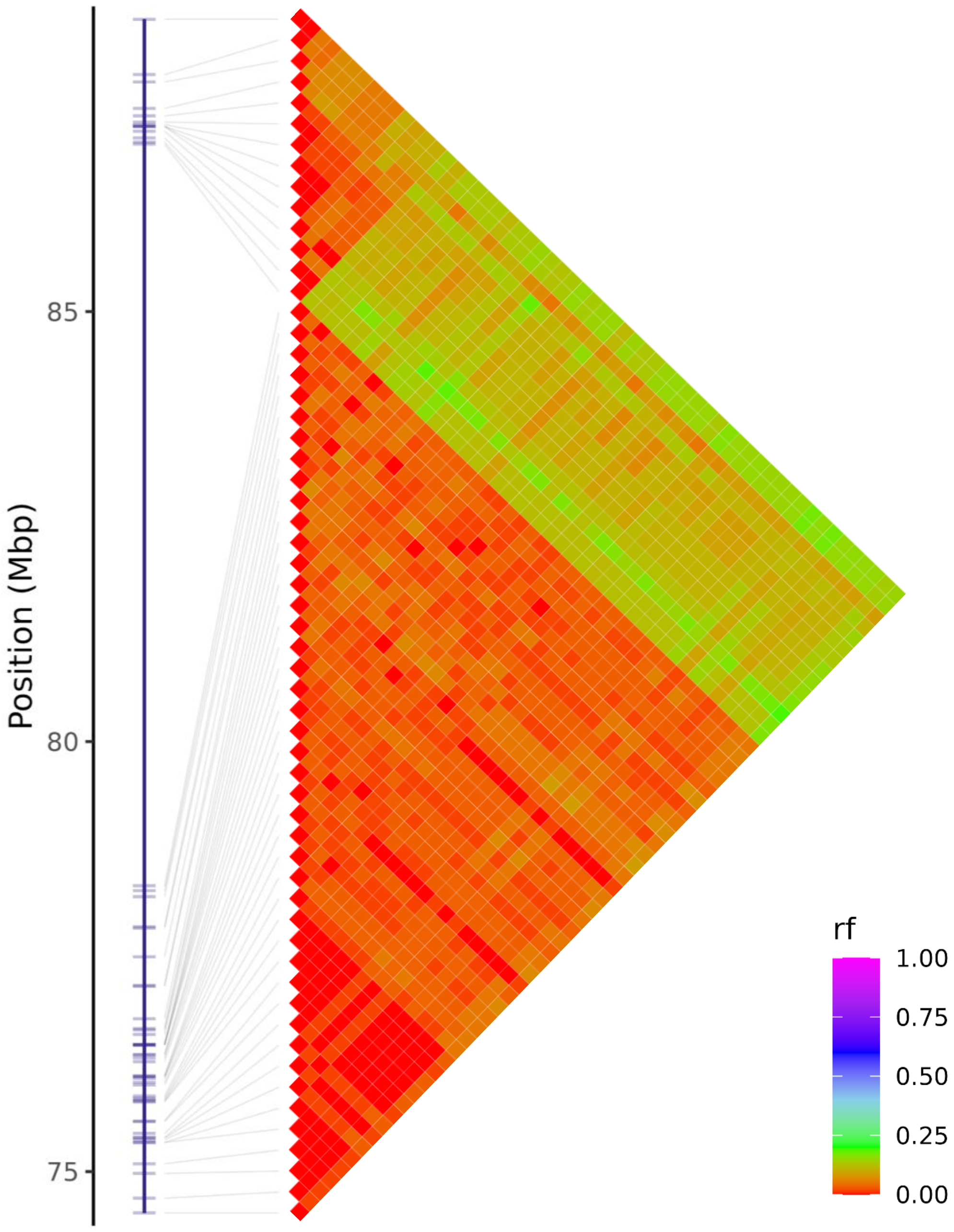


**Figure S2Q**: Linkage group 1-17 of chromosome 1, with haplotypes positioned on the monoploid reference (Chr1A) along with heatmaps of pairwise recombination fractions (rf) estimated between haplotypes.

# 1-18

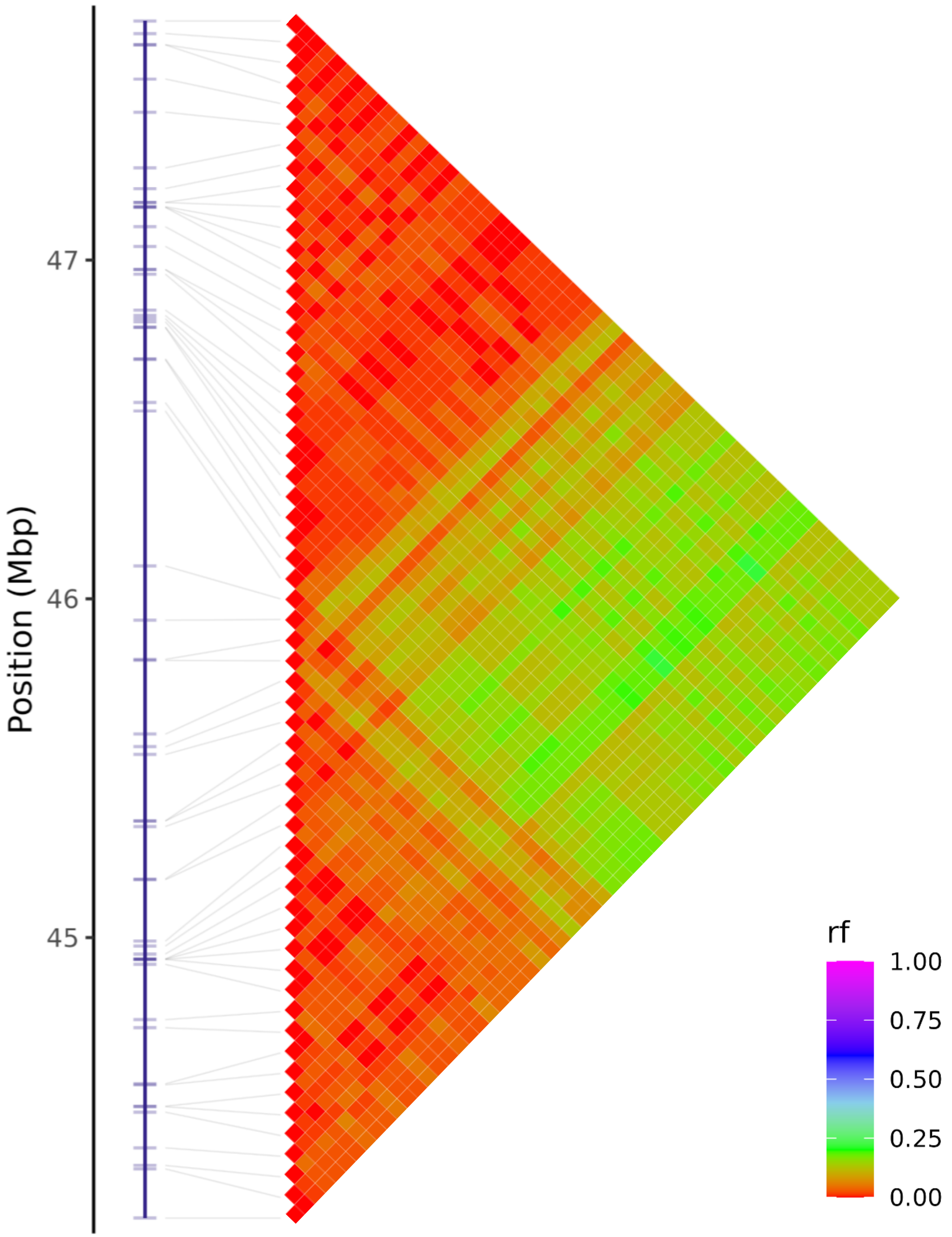


**Figure S2R**: Linkage group 1-18 of chromosome 1, with haplotypes positioned on the monoploid reference (Chr1A) along with heatmaps of pairwise recombination fractions (rf) estimated between haplotypes.
