## Supplementary material for "HaploCharmer: a Snakemake workflow for read-scale haplotype calling adapted to polyploids": Figure S3

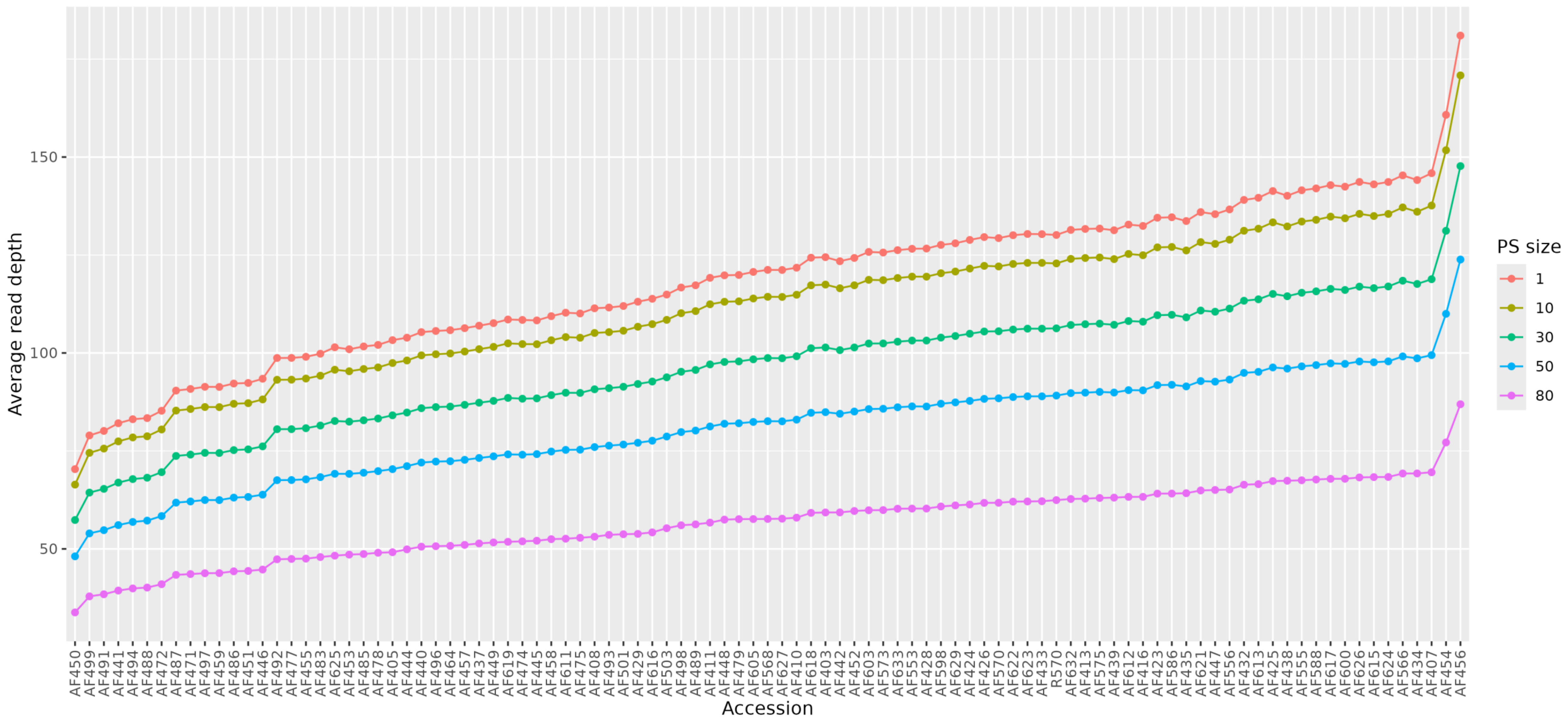


**Figure S3**: Average read depth of R570 and its self-progeny over the 86,681 PSs with varying size: 1, 10, 30, 50 or 80 bp. For each PS, only sequencing reads that fully span the entire region are considered.
