## Supplementary material for "HaploCharmer: a Snakemake workflow for read-scale haplotype calling adapted to polyploids": Figure S4

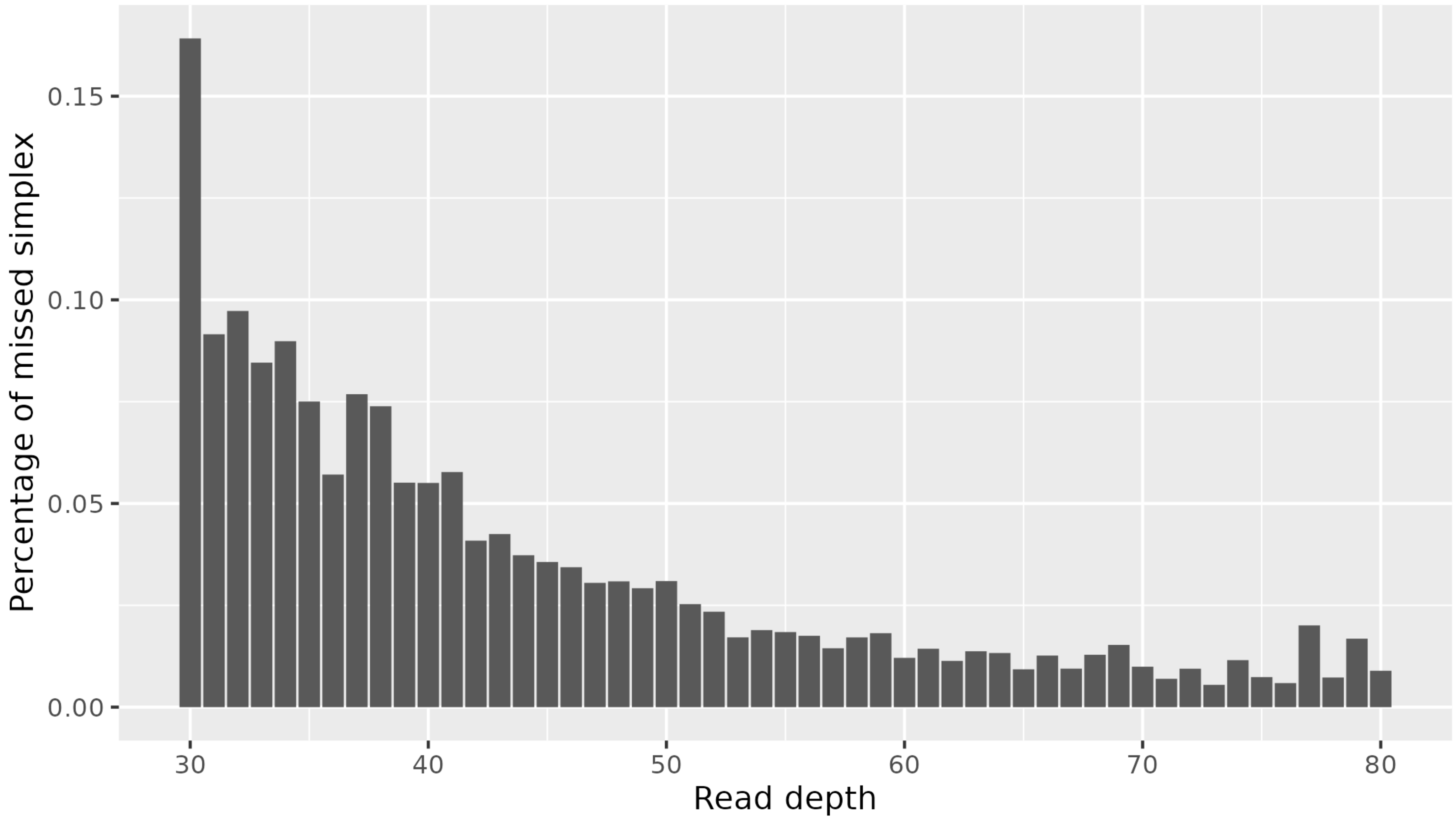


**Figure S4**: Percentage of missed single-dose haplotypes (simplex) according to read depth of the genotypic datapoint in R570. Read depths categories superior to 80 with less than 200 haplotypes were not displayed.
